# PRO-IP-seq Tracks Molecular Modifications of Engaged Pol II Complexes at Nucleotide Resolution

**DOI:** 10.1101/2023.02.04.527107

**Authors:** Anniina Vihervaara, Philip Versluis, John T. Lis

## Abstract

RNA Polymerase II (Pol II) is a multi-subunit complex that undergoes covalent modifications as transcription proceeds through genes and enhancers. Rate-limiting steps of transcription control Pol II recruitment, site and degree of initiation, pausing duration, productive elongation, nascent transcript processing, transcription termination, and Pol II recycling. Here, we developed Precision Run-On coupled to Immuno-Precipitation sequencing (PRO-IP-seq) and tracked phosphorylation of Pol II C-terminal domain (CTD) at nucleotide-resolution. We uncovered precise positional control of Pol II CTD phosphorylation as transcription proceeds from the initiating nucleotide, through early and late promoter-proximal pause, and into productive elongation. Pol II CTD was predominantly unphosphorylated in the early pause-region, whereas serine-2- and serine-5-phosphorylations occurred preferentially in the later pause-region. Serine-7-phosphorylation dominated after the pause-release in a region where Pol II accelerates to its full elongational speed. Interestingly, tracking transcription upon heat-induced reprogramming demonstrated that Pol II with phosphorylated CTD remains paused on heat-repressed genes.

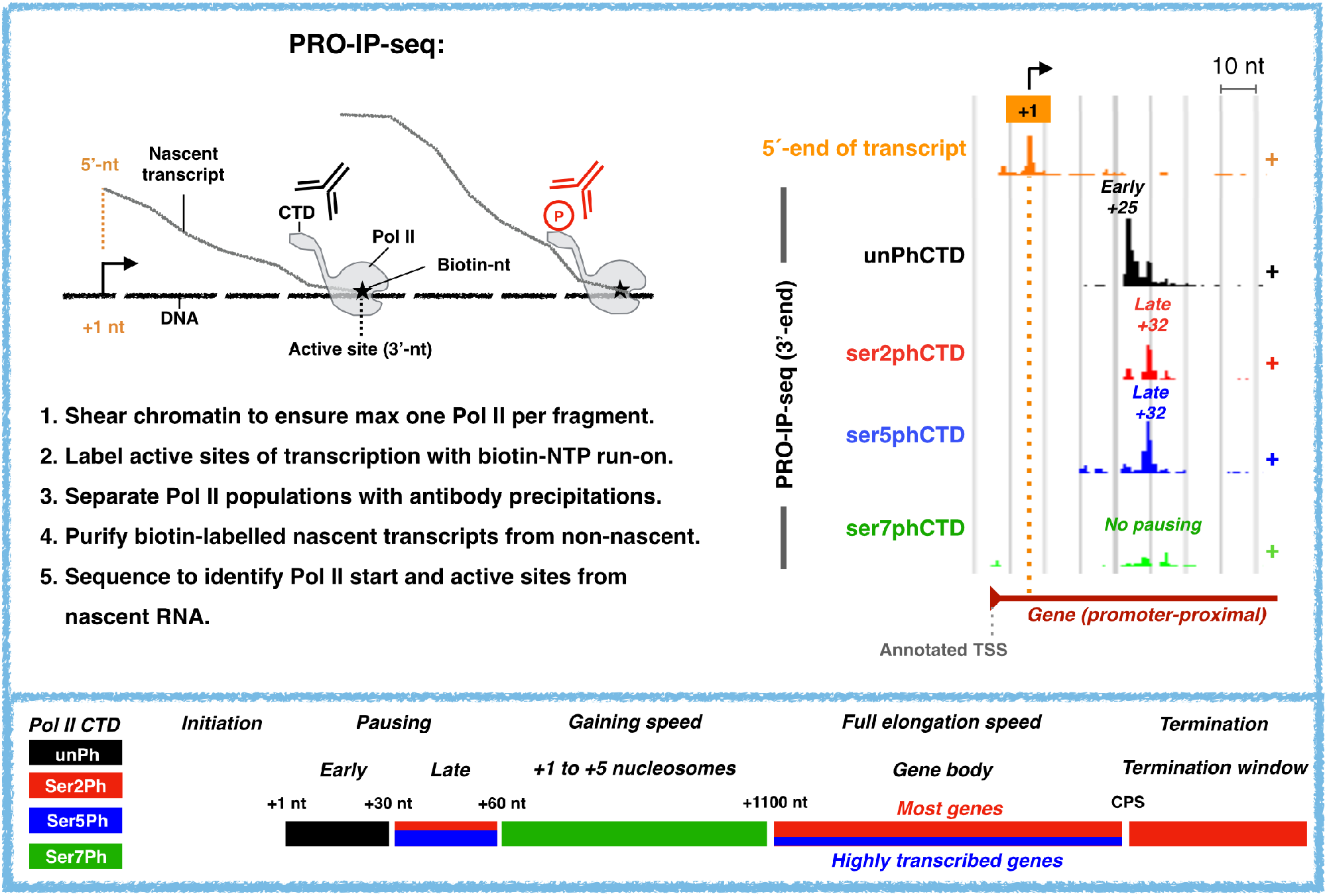

## Introduction

Regulation of transcription defines cell type-specific gene expression programs and coordinates cellular responses in health and disease. In each cell type, a distinct set of genes and enhancers are transcribed by RNA Polymerase II (Pol II) complexes. To complete a full cycle of transcription, chromatin and Pol II must undergo a series of steps: i) chromatin opening, ii) assembly of the Pre-Initiation Complex (PIC), iii) initiation of transcription, iv) promoter-proximal Pol II pausing, v) promoter-proximal Pol II pause-release, vi) productive elongation, vii) co-transcriptional RNA processing, viii) transcript cleavage, ix) transcription termination, and x) recycling of Pol II (reviewed in Wissink *et al*., 2019). These steps of transcription can be modulated by the action of transcription and elongation factors, chromatin remodelers and other co-factors, architectural proteins, and RNA-binding proteins (reviewed in Voss and Hager, 2014; Pope and Medzhitov, 2018). Additionally, transcription-regulating RNA-species and their associated RNA-binding factors can form ‘RNA clouds’ over clusters of active genes and their distal regulatory elements (reviewed in Li and Fu, 2019).

As Pol II proceeds through the transcription cycle, its phosphorylation state changes dramatically. In particular, the C-terminal domain (CTD) of the largest subunit in Pol II consists of Y_1_S_2_P_3_T_4_S_5_P_6_S_7_ repeats that undergo post-translational modifications (PTMs) and form a protean platform for regulatory interactions (reviewed in Zaborowska *et al*., 2016; Harlen and Churchman, 2017). The initiating Pol II is mainly unphosphorylated, but CDK7-mediated phosphorylation of serine-5 of the CTD disrupts Pol II interaction with TATA box Binding Protein (TBP) and the Mediator, allowing Pol II to escape the PIC (Lu et al 1992; Usheva *et al*., 1992; Wong *et al*., 2014, Jeronimo and Robert, 2014; Ebmeier *et al*., 2017). After initiation, Pol II rapidly synthetizes 20-60 nucleotides (nts) of RNA, and pauses (Rougvie and Lis, 1988; Rasmussen and Lis, 1993; Core *et al*., 2008; Kwak *et al*., 2013; Tome *et al,*. 2018). The promoter-proximal Pol II pause has emerged as a major rate-limiting step (Muse *et al*., 2007; Zeitlinger *et al*., 2007; Jonkers *et al*., 2014; Shao and Zeitlinger, 2017) *via* which genome-wide transcription programs can be coordinated (Danko *et al*., 2013; Kaikkonen *et al*., 2015; Duarte *et al*., 2016; Mahat *et al*., 2016; Vihervaara *et al*., 2017, Himanen *et al*., 2022). However, the CTD-modifications and regulatory signals along the promoter-proximal pause-region remain enigmatic. Multiple laboratories have shown that the release of Pol II from the promoter-proximal pause requires Positive Transcription Elongation Factor b (P-TEFb), the CDK9 subunit of which phosphorylates Negative Elongation Factor (NELF), transcription elongation factor SPT5 and serine-2 of Pol II CTD (reviewed in Gaertner and Zeitlinger, 2014; Core and Adelman, 2019). The release of phosphorylated NELF exposes a binding site for Polymerase Associated Factor 1 (PAF1), which converts Pol II to an elongation complex (Vos *et al*., 2018a; 2018b). After pause-release, elongation factors and histone chaperones assemble on Pol II, and topoisomerase gains its full activity, preparing the transcription complex to progress through the array of nucleosomes and other obstacles including intragenic enhancers along the gene (reviewed in Berlotserkovskaya *et al*., 2004; Mavrich *et al*., 2008, Baranello *et al*., 2016; Zumer *et al*., 2021).

To track molecular modifications of engaged Pol II complexes at high specificity and nucleotide-resolution, we developed Precision Run-On coupled to Immunoprecipitation sequencing (PRO-IP-seq). The PRO-seq portion of the protocol (Kwak *et al*., 2013) labels active sites of nascent transcripts with biotinylated nucleotides, which allows affinity purification of nascent transcripts away from the non-nascent RNA that covers the chromatin, and identification of the precise genomic positions and orientations of engaged transcription complexes. The IP portion of the PRO-IP-seq protocol selects Pol II complexes based on molecular modifications or composition using native immunoprecipitation (Gilmore and Lis, 1985; Hebbes *et al*., 1988). Here, we describe the biochemical principle of the PRO-IP-seq methodology and demonstrate how it tracks regulatory changes of the engaged transcription machinery at nucleotide-resolution. Taken together: The Pol II CTD remained unphosphorylated until the early promoter-proximal pause-region. Phosphorylation on serines 2 and 5 of the CTD occurred preferentially at the late pause-region, between +31 nt to +60 nt from the initiation. Serine-7-phosphorylation of Pol II CTD dominated from the pause-release through to the +5 nucleosome, a region where Pol II assembles elongation factors and gains speed (Jonkers *et al*, 2014). Our results confirmed serine-2-phosphorylation as the most distinctive CTD mark of active elongation; however, serine-5-phosphorylation was also detected throughout the gene’s body and correlated with higher transcriptional activity of a gene. Finally, tracking changes in Pol II CTD upon rapid transcriptional reprogramming by heat shock revealed that the CTD of paused Pol II can be, and often is, phosphorylated at serines 2 and 5.

## Results

### PRO-IP-seq tracks molecular modifications of nascent transcription complexes at nucleotide-resolution

Run-on reactions map engaged Pol II complexes at nucleotide-resolution by incorporating a single biotin-tagged nucleotide into the active sites of transcription (Kwak *et al*., 2013). To couple run-on reaction to immunoselection of Pol II populations, we performed the run-on reaction on chromatin (Chu *et al*., 2018). Chromatin was prepared from three biological replicates of non-stressed (NHS) and 30 min at 42°C heat shocked (HS30) human K562 cells and fragmented to < 80 base pairs (bp) using light sonication and DNase I (Figure 1A and Figure S1A). Since elongating Pol II footprints 43 bp of DNA *in vitro* (Saeki and Svejstrup, 2009), each fragment is expected to contain at most a single engaged Pol II complex. Even at promoter-proximal pause sites where Pol II can be at very high density, a single engaged Pol II sterically hinders new initiation until transcription has proceeded beyond the pause (Shao and Zeitlinger, 2017; Gressel *et al*., 2019). Note also that DNase I does not cleave RNA and has a low activity towards RNA:DNA hybrids (Vanecko and Laskowski, 1961; Huang *et al*., 1996), leaving the nascent transcripts intact. Thus, the assay is designed to map individual Pol II complexes at the resolution and sensitivity afforded by PRO-seq.

**Figure 1.**
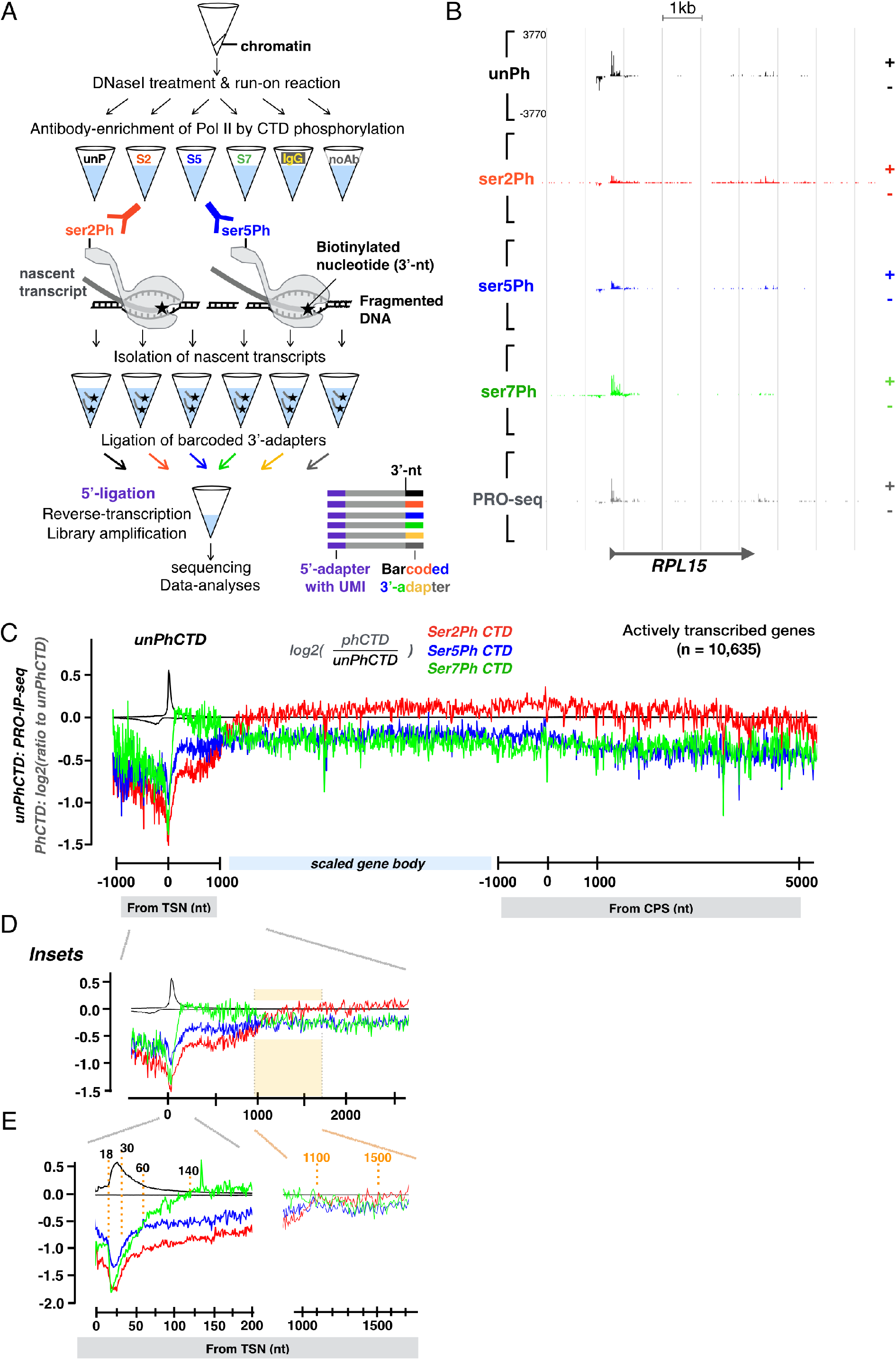
PRO-IP-seq maps molecular modifications of nascent transcription complexes at nucleotide resolution. **A)** Schematic presentation of the biochemical steps of PRO-IP-seq: Isolated chromatin is fragmented and a single biotinylated nucleotide incorporated into active sites of transcription. Transcription complexes are selected with antibodies against the C-terminal domain of Pol II (CTD), and nascent transcripts separated from non-nascent. The nascent transcripts are ligated to barcoded 3’-adaptors, and the barcoded samples pooled for equal handling. After removal of 5’-cap, UMI-containing 5’-adaptors are ligated. S2, S5 and S7 refer to phosphorylations of serine residues at Pol II CTD. B) Representative example gene, *RPL15*, showing distribution of engaged Pol II with indicated status of CTD in non-treated K562 cells. The Y-scale is linear and the same in all tracks. **C**) Comparison of distinct CTD phosphorylations relative to the unphosphorylated form of Pol II CTD across the promoter-proximal region (linear scale from −1000 to +1000 nt from transcription start nucleotide, TSN, +1), gene body (scaled to 50 bins per gene), and CPS & termination window (linear scale from −1000 to +5000 from the CPS). The CTD modifications were mapped across all actively transcribed genes in untreated human K562 cells (n=10,635). Average PRO-IP-signal is indicated for unphosphorylated Pol II CTD (black transcription profile), and the ratio of phosphorylated relative to unphophorylated CTD (red, green and blue) shown in log2 scale. D-E) Insets of the promoter-proximal region. The dotted orange lines indicate nt-resolution distance from the TSN. In **D-E** each window is 1 nt. The Y-scales are as in **C**).

The run-on reaction was conducted in the presence of whole-genome spike-in derived from mouse cells, and the run-on chromatin divided into distinct tubes for pull-downs with monoclonal antibodies against the Pol II CTD (Figures 1A and S1B). As a negative control, we used non-specific antibody (IgG), and as a positive control, we examined total nascent (PRO-seq) RNAs (noAb). In this strategy, the same chromatin and labelled nascent RNAs served as starting material for all antibody pulldowns and controls in a condition. After the immunoprecipitation, the nascent transcripts were purified and barcoded, and samples re-pooled to ensure identical handling (Figure 1A). The size-distribution of reverse-transcribed nascent RNAs (Figure S1B) was as expected for nascent RNA-seq libraries (Mahat *et al*., 2016a), and the Pol II CTD pulldowns generated nucleotide-resolution density profiles characteristic of nascent transcription (Figure S1B-C). The data was initially normalized and examined with Reads Per Million of mapped reads (RPM) strategy (Figure S2A). To refine the normalization strategy, we created a normalization factor corrected control RPM (nf-cRPM) that does not assume a similar total transcription between the samples (see online Materials and Methods). The nf-cRPM yielded highly similar density profiles between PRO-IP-seq replicates, recapitulated known patterns of gene and enhancer transcription, brought distinct run-on experiments to comparable y-scale, and generated equivalent PRO-IP-seq profiles with different antibodies against the same Pol II CTD modification (Figures S1B, S2B-D).

### PRO-IP-seq identifies +1 nucleotides from nascent transcripts

Tracking nucleotide-resolution changes in transcription from initiation through the promoter-proximal pause requires precise identification of the +1 nucleotide (nt) and the active sites of transcription. The Transcription Start Sites (TSSs) derived from the reference genomes generally report the longest form of each mRNA isoform, which does not necessarily coincide with the most prominent nucleotide that initiates the transcription. To precisely map the Transcription Start Nucleotide (TSN, +1 nt) at each gene, we used the 5’-ends of nascent RNAs that associate with Pol II in our PRO-IP-seq data (Figure S3A-D). The 5’-ends of PRO-IP-seq reads enriched virtually at the same +1 nt regardless of the CTD modification (Figure S3A-C), which supports the concept that the CTD phosphorylation does not affect the site of initiation, and also that sites of initiation do not affect CTD phosphorylation patterns. In agreement, the same +1 nt was detected in distinct PRO-seq datasets that do not perform an antibody pulldown (Figure S3A-C). Worth noting is that the TSN identified from 5’-ends of nascent transcripts resided on average 77 nt (median 55 nt) downstream of the reference genome annotated TSS (Figures S3B and S3D). The discrepancy between reference genome annotations and sequencing of 5’ ends of RNAs has been previously noted (Nechaev *et al*., 2010; Tome *et al*., 2018). These and our current study highlight the importance of identifying the exact +1 nt from nascent transcripts in the cells being studied to provide a correct reference point to map chromatin architecture and the precise genomic positions of the transcription machinery.

### Genes are divided into five domains of Pol II CTD modifications

Visualizing Pol II CTD modifications across individual genes uncovered high enrichment of Pol II with unphosphorylated CTD at the promoter-proximal pause region (Figure 1B). Serine-2-phosphorylated CTD, instead, was distributed over gene bodies, and serine-7-phosphorylated CTD occupied early coding regions (Figure 1B). To broadly investigate changes in Pol II CTD phosphorylation across genes, we graphed the ratio of phosphorylated CTD against unphosphorylated CTD at all transcribed genes (Figure 1C-E). These analyses uncovered five domains of Pol II CTD modifications (Figure 1C): 1) At the early pause-region, the CTD was primarily unphosphorylated, while 2) in the late pause region, the CTD showed rapidly increasing phosphorylation on serines 2, 5, and 7. In the 3) immediate region downstream of the pause, the CTD contained strikingly high serine-7-phosphorylation, which was replaced by 4) high serine-2-phosphorylation along the region from 1.2 kb from initiation to the cleavage and polyadenylation site (CPS). Finally, 5) in the termination window, the CTD retained serine-2-phosphorylation but lost serine-5-phosphorylation. Changes in Pol II CTD occurred over short genomic distances, as demonstrated by the steep incline in serine-7-phosphorylation over promoter-proximal pause-release, and the prominent shift from serine-7-to serine-2-phosphorylation at +1000 to +1200 from the TSN (Figure 1D).

The 52 heptad repeats in human Pol II can undergo multiple types of post-translational modifications (reviewed in Zaborowska *et al*., 2016). Consequently, the availability of unphosphorylated CTD could be affected by modifications beyond serine-2, serine-5, and serine-7 phosphorylation. To ensure appropriate comparison of Pol II CTD state, we also analyzed the Pol II CTD modifications relative to the total level of transcription (PRO-seq only, noAb). These analyses showed highly similar patterns and densities of Pol II CTD modifications as detected when comparing phosphorylations relative to unphosphorylated CTD (Figure S4). Taken together, nucleotide-resolution PRO-IP-seq shows Pol II to initiate with unphosphorylated CTD, to become increasingly phosphorylated at serines 2, 5 and 7 as it progresses over the promoter-proximal pause region, to then be highly enriched for serine-7 phosphorylation from the point of pause-release to +1.2 kb from the initiation. After +1.2 kb from the initiation Pol II CTD becomes abundantly phosphorylated at serine-2 over the gene body and termination window, which is accompanied by reduced levels of serine-5-phosphorylation after the CPS (Figures 1B-D and S4).

### Pol II CTD is primarily unphosphorylated until +30nt from the transcription start nucleotide

Early targeted-gene studies showed genes contain two distinct promoter-proximal pause peaks within a single pause-region (Rasmussen and Lis, 1993). Indeed, coordinated, genome-wide mapping of initiation and pausing revealed two distinct enrichments of engaged Pol II at the promoter-proximal region: an early pause until +30 nt, and a late pause between +31 to +60 nt, from the TSN (Tome *et al*., 2018). Here, tracking CTD modifications of engaged Pol II at nucleotide-resolution uncovered that unphosphorylated CTD dominates from the initiation until the early promoter-proximal pause-region (Figure 1). Indeed, unphosphorylated Pol II was most enriched at +25 nt from the TSN, after which, the relative levels of serine-2-, serine-5- and serine-7-phosphorylations increased (Figure 1E). In accordance, Pol II with unphosphorylated CTD often occupied the pause closer to the TSN, while Pol II with serine-2- and serine-5-phosphorylated CTD was enriched at pause sites more downstream (Figure 2A). At highly transcribed genes, such as *betaactin (ACTB*), unphosphorylated CTD was the primary form of paused Pol II detected across the pause region (Figure 2B). Mapping the most prominent pause coordinate of each Pol II CTD modification confirmed that a clear pause signal was most often found with an antibody to the unphosphorylated CTD epitope (77% of all active genes). In contrast, pausing of Pol II bearing phosphorylated CTD was detected at 59% (serine-2 and serine-5), or 56% (serine-7) of transcribed genes (Figure 2C). Furthermore, unphosphorylated CTD enriched primarily at the early pause-region, peaking between +18 to +30 nt from the TSN (Figure 2C). Pol II with serine-2-, serine-5- or serine-7-phosphorylated CTD were, instead, more evenly distributed over the pause regions, showing a preference toward the late pause (Figure 2C).

**Figure 2.**
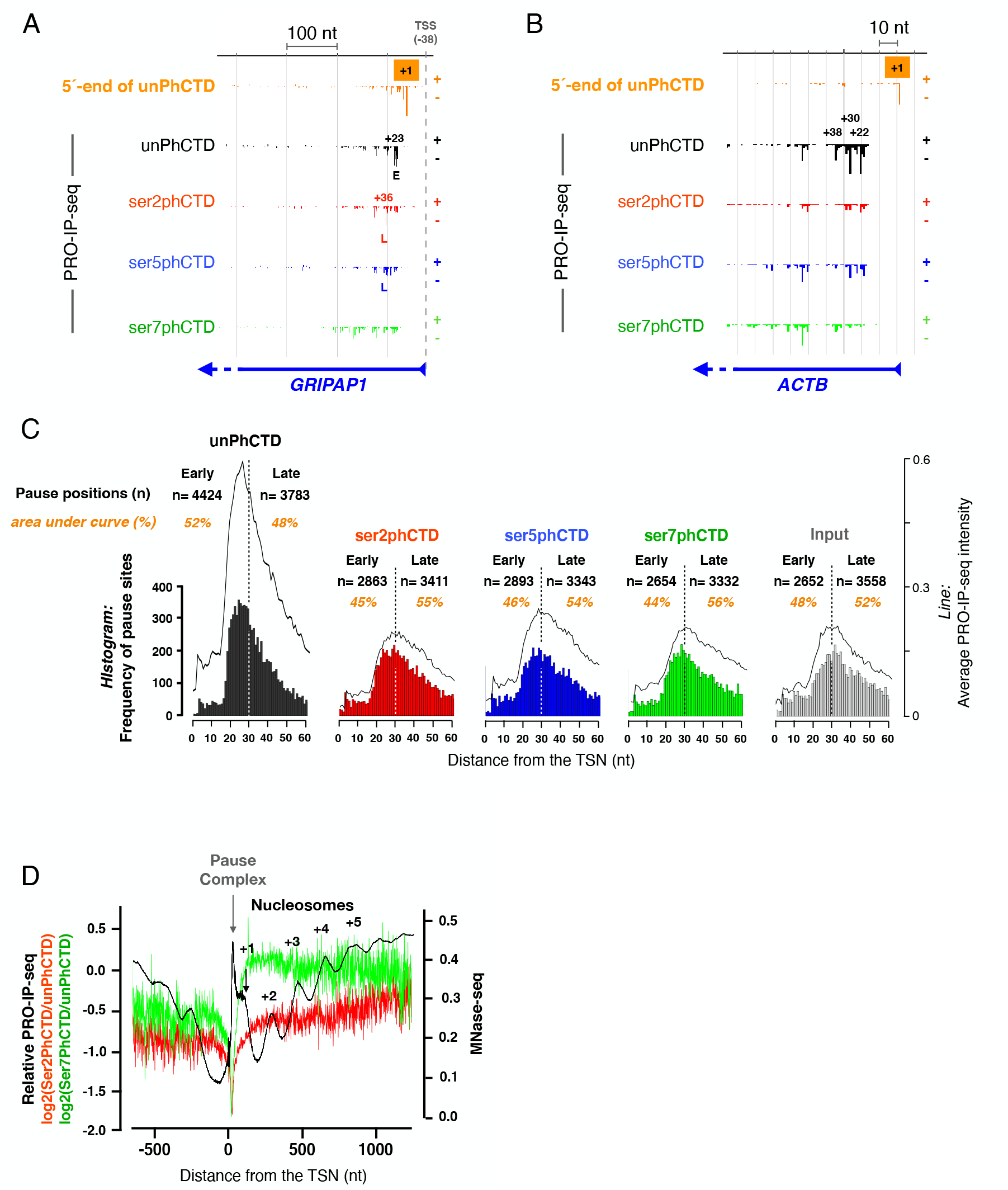
Pol II with unphosphorylated CTD is enriched at the early pause. **A-B)** PRO-IP-seq identifies transcription start nucleotide (TSN, +1) and the exact pause sites of engaged Pol II complexes, as exemplified at *GRIPAP1* **(A)** and *ACTB* **(B)** promoters. At *GRIPAP1* Pol II with unphosphorylated CTD is enriched at early pause (E), while Pol II with serine-2-posphorylated CTD localises to the late (L) pause. At *ACTB* un phos phorylated CTD has the strongest enrichment on all the pause sites. The TSN is identified using 5’-ends of PRO-IP-seq reads from the unPhCTD. The numbers indicate genomic distance (nt) from the TSN. The RefSeq-annotated transcription start site (TSS, dashed grey line) is depicted at *GRIPAP1* and its distance to the TSN indicated. **C)**. Histograms showing the distance of the strongest pause coordinate to the TSN. The count (n) of pause sites detected at early (+1 to +30 from the TSN) and late (+31 to +60 from TSN) pause-region are indicated. The line above the histogram shows Pol II density across the pause-region. The percentage of Pol II at the early and late pause are counted as areas under the curve. **D)** Relative levels of serine-2- and serine-7-phosphorylated CTD overlaid on MNase-seq detected nucleosomes at the open chromatin region of gene promoters. The PRO-IP-seq and MNase-seq were quantified on all transcribed genes in human K562 cells (n=10,635). The MNase-seq data is from ENCODE (Consortium EP, 2011)

### Serine-7-phosphorylation of Pol II CTD dominates from the pause-release until +5 nucleosome

Pol II CTD had low levels of serine-7-phosphorylation at the early pause-region (Figure 1E). Moreover, serine-7-phosphorylated Pol II did not show prominent enrichment at either early or late pause site (Figure 2A-B). Instead, phosphorylation at serine-7 rapidly increased after the promoter-proximal pause, becoming the most prominent CTD phosphorylation after +60 nt from the TSN, and reaching its highest levels in +140 to +160 nt window (Figure 1E). In this early coding region, Pol II assembles elongation factors, gains speed, and encounters the +1 nucleosome (reviewed in Jonkers and Lis, 2015). To investigate Pol II CTD phosphorylation with respect to the nucleosomes, we overlayed MNase-seq to the relative enrichments of serine-2- and serine-7-phosphorylations (Figure 2D), centering the signal to the TSN. These analyses uncovered serine-7-phosphorylation to peak at the +1 nucleosome and remain high until +5 nucleosome at the end of the exonuclease-accessible region (Figure 2D). Serine-2-phosphorylation, instead, increased through the ordered array of +1 to +5 nucleosomes becoming the dominant Pol II CTD modification at the end of the MNase-accessible region (Figures 1E and 2D).

### Actively transcribed gene bodies have high levels of serine-2 and serine-5-phosphorylated CTD

At an average gene, serine-2-phosphorylation was the most dominant CTD modification from +1200 nt from the TSN to the end of the termination window (Figure 1B-E). However, many highly transcribed genes had prominent levels of serine-5-phophorylation across the gene body (Figure 3A). To investigate whether serine-5-phosphorylation co-occurred with high gene activity, we grouped genes based on productive elongation (Figures 3C and S5A-B). While genes with low and moderate activity were covered by serine-2-phosphorylated CTD along the gene body (Figure 3B-C), genes with the highest transcriptional activity contained comparable levels of serine-2- and serine-5-phosphorylation at the CTD (Figure 3A, C). In contrast, the CPS and termination window were dominated by serine-2-phosophorylated CTD in every gene group studied (Figure 3A-C).

**Figure 3.**
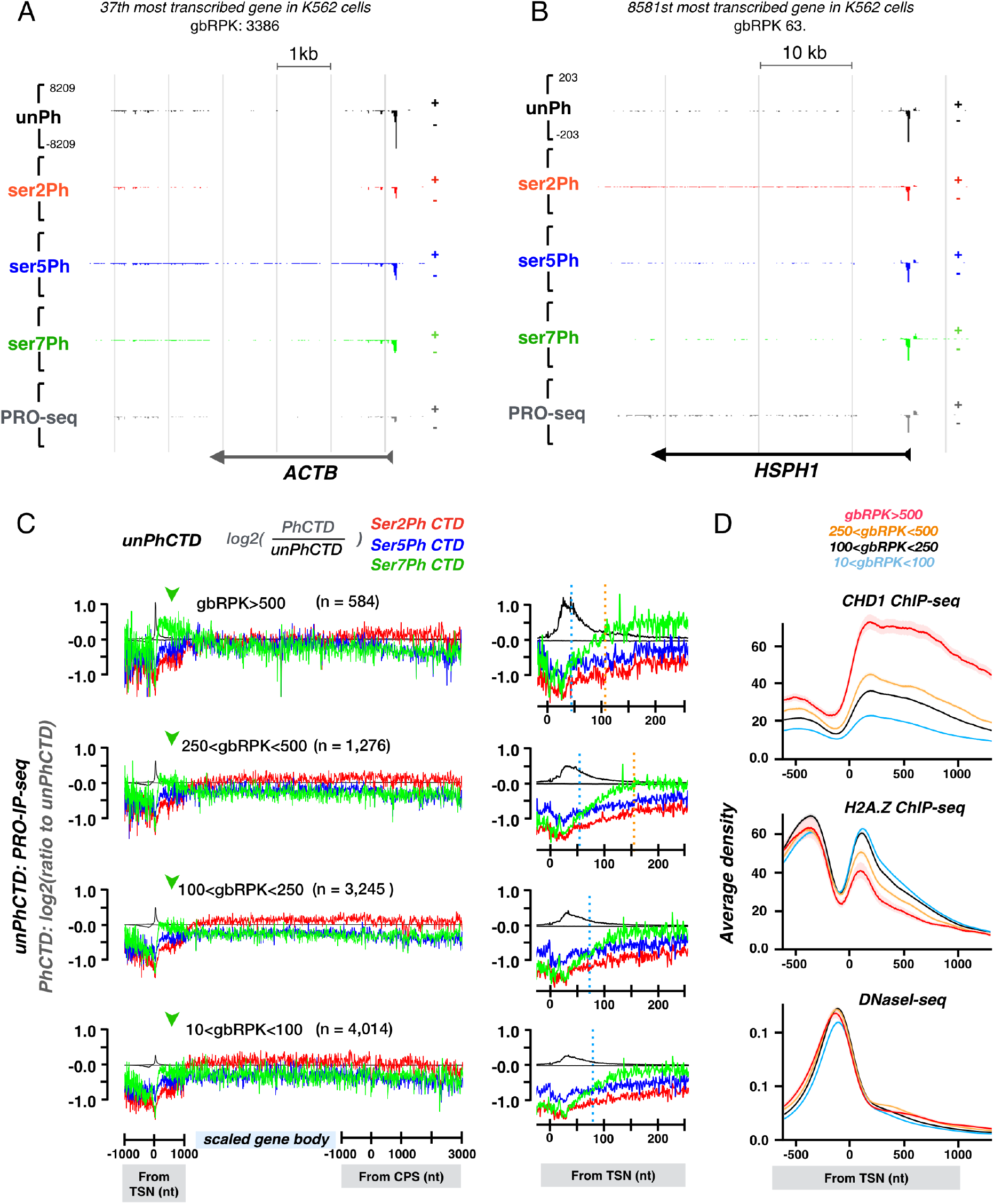
Transcriptional activity is reflected in the CTD code. **A)** *Beta-actin (ACTB*), one of the most active genes in human, shows 1) prominent pausing of Pol II, primarily with unphophorylated CTD, 2) ser-7-phosphorylation extending to early coding region, and 3) prominent ser-5-phosphorylation of the CTD across the gene body. **B**) *HSPH1*, a lowly transcribed gene in nonstressed human cells, shows 1) high Pol II pausing that is detected with all the investigated antibodies, 2) no enrichment of ser-7-phosphorylation at the early coding region, and 3) ser-2-phosphorylation dominating across the gene body. **C**) Genes were grouped based on their transcriptional activity (reads per kilo bases of gene body, gbRPK) and the CTD phosphorylations compared against the unphospharylated form of the CTD. Promoter-proximal region: linear scale from −1000 to +1000 nt from transcription start nucleotide (TSN). Gene body: scaled to 50 bins per gene. CPS& termination window: linear scale from −1000 to +5000 from the CPS. Average density of unphosphorylated Pol II CTD is indicated as black transcription profile, and the ratio of phosphorylated per unphophorylated CTD (red, green and blue) shown in log2 scale. The insets show the early coding region. The dotted blue vertical line indicates where the relative amount of ser-7-phosphorylation exceeds the relative amount of ser-5-phosphorylation. The dotted orange vertical line indicates where the relative amount of ser-7-phosphorylation exceed the relative amount of un-phosphorylated CTD. **D**) CHD1 ChlP-seq, H2A.Z ChlP-seq and DNasel-seq signal (Consortium EP, 2011) at the promoter-proximal region of indicated genes. The shaded area indicates 12.5-87.5% interval for each group.

### Serine-7-phosphorylation at the early coding region predicts productive elongation along the gene

Analyzing Pol II CTD modifications with respect to transcriptional activity uncovered serine-7-phosphorylation at the early coding region to predict the level of productive elongation downstream at the gene body (Figures 3C and S5A-B). The more productive elongation the gene had, the earlier the phosphorylation of serine-7 occurred and the higher its relative enrichment became (Figure 3D). As comparison, relative levels of serine-2 and serine-5-phosphorylations were similar across the early coding regions regardless of the gene’s transcriptional activity (Figure 3C). Next, we overlaid MNase-seq and histone 4 acetylation (H4ac) ChIP-seq with the PRO-IP-seq signals of distinct gene groups. The positioning of +1 to +5 nucleosomes was comparable regardless of the transcriptional activity of the gene (Figure S5C), but their active histone marks, indicated as H4ac levels (Vettese-Dadey *et al*., 1996; Müeller *et al*., 2017; Furukawa *et al*., 2020), increased with the productive elongation through the gene (Figure S4D). Increased chromatin activity at serine-7-rich regions was further supported by the positive correlation of Chromodomain Helicase DNA binding protein 1 (CHD1), and negative correlation of H2A.Z (Figure 3D). Of note is that the accessibility of the promoter and the early coding region to DNase I was similar regardless of the gene’s transcriptional activity (Figure 3D). These results suggest that serine-7-phosphorylation at the CTD is involved in establishing the processivity of Pol II through the gene. This processivity is gained at the early coding region where elongation factors assemble on Pol II and the transcription elongation rapidly accelerates (Jonkers *et al*., 2014). Indeed, the study of Pol II elongation rates by Jonkers *et al*.showed that Pol II undergoes its major acceleration during the first 1-2kb. In accordance, once Pol II had proceeded beyond +1500 nt from the TSN, the levels of serine-phosphorylations did not change appreciably along the gene, either when analyzed as average enrichments over unphosphorylated CTD or from individual genes (Figures 1B-C, 3A-C, S1D, S2).

### Heat shock reorganizes Pol II CTD phosphorylation across the genome

Heat shock triggers an instant reprogramming of transcription (Duarte *et al*., 2016; Mahat *et al*.,2016; Vihervaara *et al*., 2017; 2021). This rapid transcriptional response is primarily coordinated at the rate-limiting step of Pol II pause-release (reviewed in Vihervaara *et al*., 2018): At thousands of heat-repressed genes, Pol II accumulates at the promoter-proximal region. In striking contrast, heat-activated genes overcome the global repression of pause-release by recruiting potent *trans*activators that trigger the release of paused Pol II into elongation. The master *trans*-activator at heat-activated genes is Heat Shock Factor 1 (HSF1) that can *trans*-activate chaperone, cochaperone, and polyubiquitin genes (Hahn *et al*., 2004; Mendillo *et al*., 2012; Vihervaara *et al*., 2013; reviewed in Vihervaara *et al*., 2014) both *via* proximal and distal regulatory elements (Ray *et al*.,2019; Vihervaara *et al*., 2021; Himanen *et al*., 2022). Here, we used heat shock to track Pol II CTD modifications at 1) heat-induced genes where paused Pol II is efficiently released into elongation, and 2) heat-repressed genes where the release of paused Pol II is inhibited.

Heat shock induced profound changes in the phosphorylation status of Pol II CTD, both at individual genes (Figure S6A-B) and across large regions within topologically associated domains (TADs; Figures 4A and S6C). Intriguingly, Hi-C maps showed TADs with highly heat-induced genes to contact TADs with clusters of heat-repressed genes (Figures 4A and S6). These connected primary TADs resided in larger nested TADs as exemplified with two major heat-induced loci, HSP70 and HSP90, connecting to a strongly heat-repressed histone locus (Figures 4A-B). Histone genes in all three major clusters showed Hi-C and ChIA-PET identified connections to *HSP70* and *HSP90* genes (Figure 4B-C). At the histone locus, all Pol II CTD modifications were reduced upon stress, and the transcription machinery concentrated at the promoter-proximal pause (Figure 4D). Concordantly, the HSP loci gained a massive transcriptional activation involving all Pol II CTD modifications (Figure 4D). The kinetic link between HSP induction and histone repression has been previously shown in *Drosophila* (O’Brien and Lis, 1993), mouse (Mahat 2016) and human cells (Vihervaara, 2017). Together, these data indicate that genes with similar responses are separated into their own sub-TADs (Figure S6), but connections between *loci* might facilitate the diffusion of Pol II and its regulators from heat-repressed to heat-induced genes.

**Figure 4.**
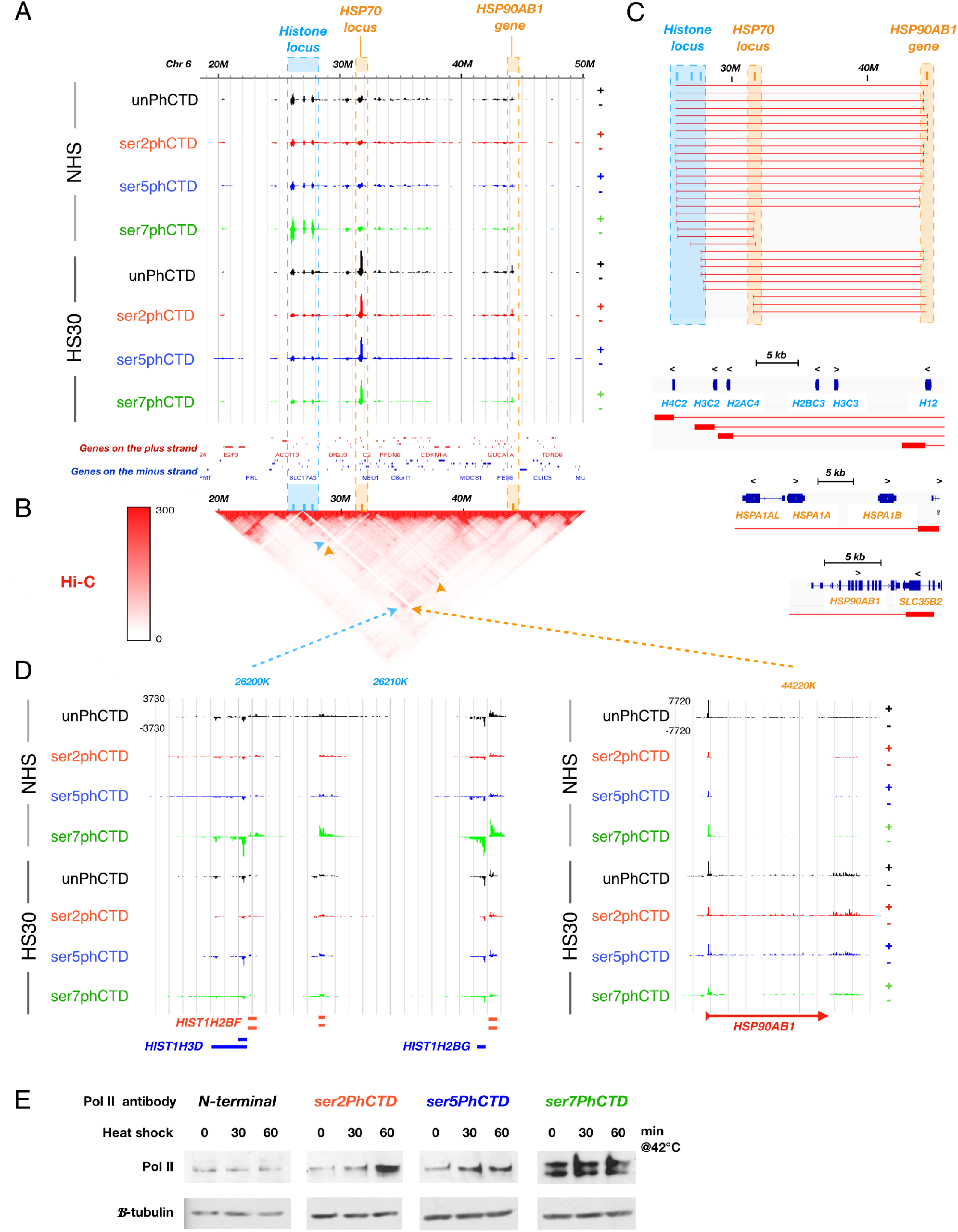
Heat shock reprograms Pol II CTD phosphorylation. **A)** A 30-Mb region of chromosome 6, showing positions of *histone, HSP7O and HSP9O loci* and changes in Pol II CTD phosphorylation B) Hi-C map indicating connections between
*HSP* and *histone loci* within a nested TAD. **C**) ChlA-PET-identified connections (red blocks and lines) that occur between *histone, HSPA1A-HSPA1B* and *HSP90AB1* genes. Expanded browser profiles below the connection data exemplify connections originating from the *histone, HSPA1A-HSPA1B*, and *HSP90AB1 loci*. **D)** Part of the histone locus (left panel) and the *HSP90AB1* gene (right panel) showing changes in Pol II CTD. E) Levels of total Pol II and CTD phosphorylations upon heat shock.

### Heat shock causes a global increase in serine-5-phosphorylated CTD

Heat shock is known to increase the proportion of phosphorylated, high-molecular weight, Pol II (Dubois *et al*., 1991; 1997). We found that the total cellular pool of Pol II gradually gained phosphorylation at serine-5 of the CTD during heat stress (Figures 4E and S7A). Also, serine-2-phosphorylation increased during heat shock, but the increase was mainly detected after 30-minutes of heat stress (Figures 4E and S7A). Tracking phosphorylation of Pol II across genome showed heat-induced genes to contain high levels of both serine-2- and serine-5-phosphorylated CTD (Figure S6A,S7B). Even at heat-repressed genes, the Pol II that did enter productive elongation was enriched with serine-5 phosphorylation of the CTD (Figure S6B,S7C).

### Pol II CTD gains serine 2 and serine 5 phosphorylation at the pause-region of heat-induced genes

Heat shock increases recruitment of Pol II to heat-activated genes (Mahat *et al*., 2016; Vihervaara *et al*., 2017). Interestingly, the promoter-proximal regions of heat-activated genes gained primarily Pol II bearing serine-2- and serine-5-phosphorylated CTD (Figure 5A). Moreover, the unphosphorylated Pol II shifted from the early to the late pause (Figure 5B-C). In comparison, serine-7-phosphorylation only modestly increased along the gene, and Pol II with serine-7-phosphorylated CTD did not appear to pause at the promoter-proximal region (Figure 5A-C). These results support our nucleotide-resolution profiles in the NHS condition (Figures 1–3), indicating that phosphorylation of serine 7 at the CTD occurs after Pol II has been released from the promoter-proximal pause into early coding region.

**Figure 5.**
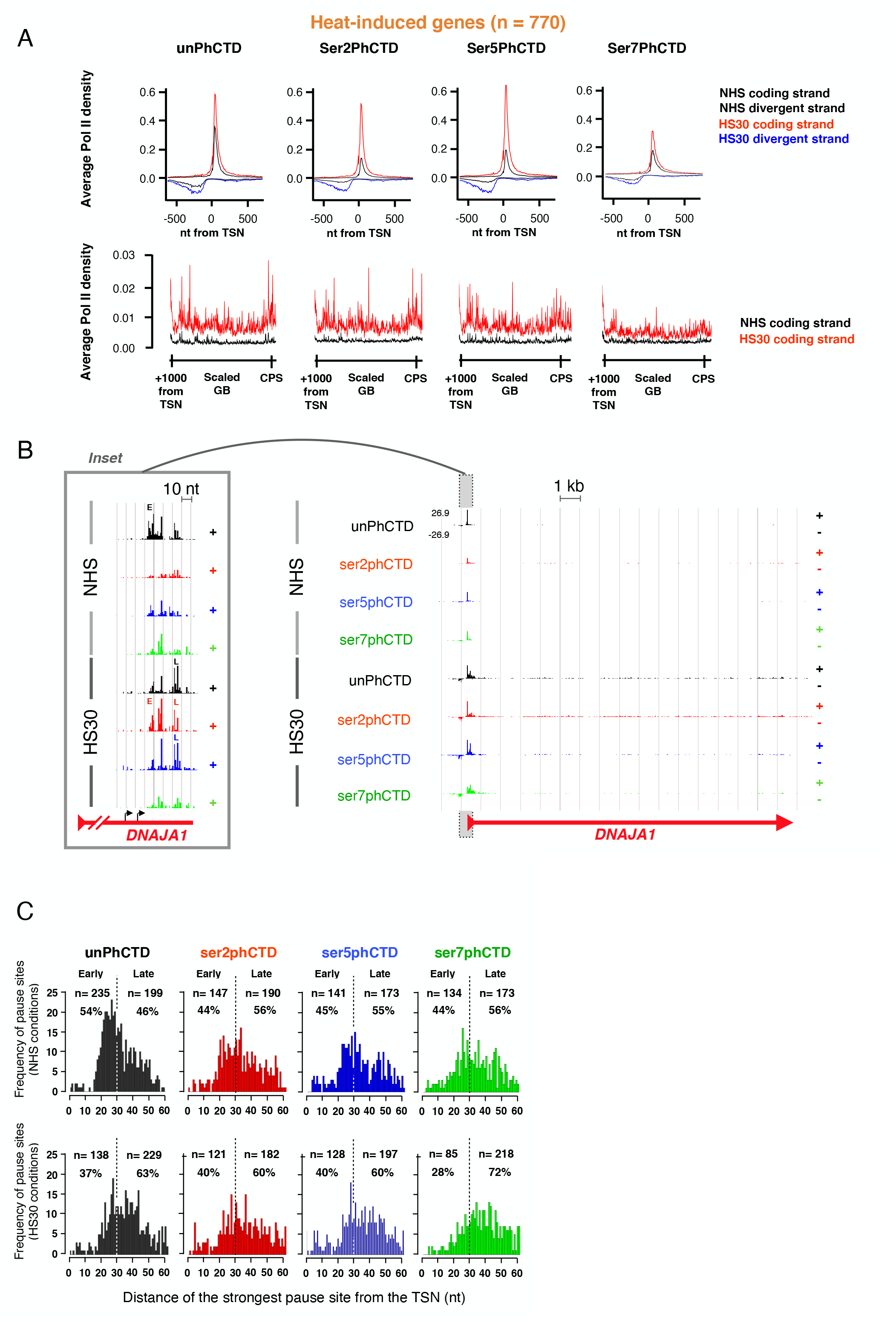
Heat-activation launches Pol II into elongation and concentrates promoter-proximal Pol II to distal pause. **A)** Average PRO-IP-seq densities with indicated Pol II CTD modification at promoter-proximal regions (upper panels) and gene bodies (lower panels) of heat-activated genes (n=770). The black lines depict PRO-IP-seq signal in non-stressed (NHS) and red (coding strand) and blue (non-coding strand) in 30 minute heat shocked (HS30) cells.) TSN; Transcription start nucleotide. **B**) Distribution of CTD modifications on *DNAJA1* gene showing heat-induced wave of Pol II with all detected CTD marks. The inset shows promoter-proximal region, depicting the distribution of CTD modifications to early (E) and late (L) pauses. The black arrows indicated two distinct +1 nucleotides (transcription initiation nucleotides, TINs) detected from the nascent transcripts. **C**) Histograms depicting the coordinate of the highest pause on each promoter-proximal region of heat-induced gene in NHS (upper panels) and upon HS30 (lower panels). The count (n) and fraction (%) of pauses detected on early (+1 to +30 from the TSN) and late (+31 to +60 from TSN) are indicated.

### Pol II with phosphorylated CTD remains paused at heat-repressed genes

Next, we investigated how the phosphorylation status of CTD changes at repressed genes where heat shock prevents the entry of Pol II into productive elongation. Strikingly, inhibiting the pause-release caused accumulation of Pol II with serine-2 and serine-5 phosphorylated CTD at the early and late pause regions of heat-repressed genes (Figures 6A-B). The pausing of Pol II with phosphorylated CTD was particularly evident at long genes (Figure 6C); Pol II molecules that were downstream of the pause when the heat stress began continued transcribing, causing a wave of clearing Pol II molecules from the coding regions (Figure 6C). These clearing Pol II molecules maintained their phosphorylation status. In contrast, levels of Pol II at the pause-region increased during heat shock and the pattern of CTD phosphorylation changed. Pol II with unphosphorylated CTD epitope was enriched in the early pause in NHS condition, whereas Pol II with serine-2 and serine-5-phosphorylated CTD epitope occupied the late pause (Figure 6C). Upon heat shock, the pause-release was inhibited and Pol II bearing unphosphorylated, serine-2-phosphorylated and serine-5-phosphorylated CTD concentrated to the early and late pauses (Figure 6C). These results demonstrate that Pol II with serine-2- and serine-5 phosphorylated CTD can remain paused.

**Figure 6.**
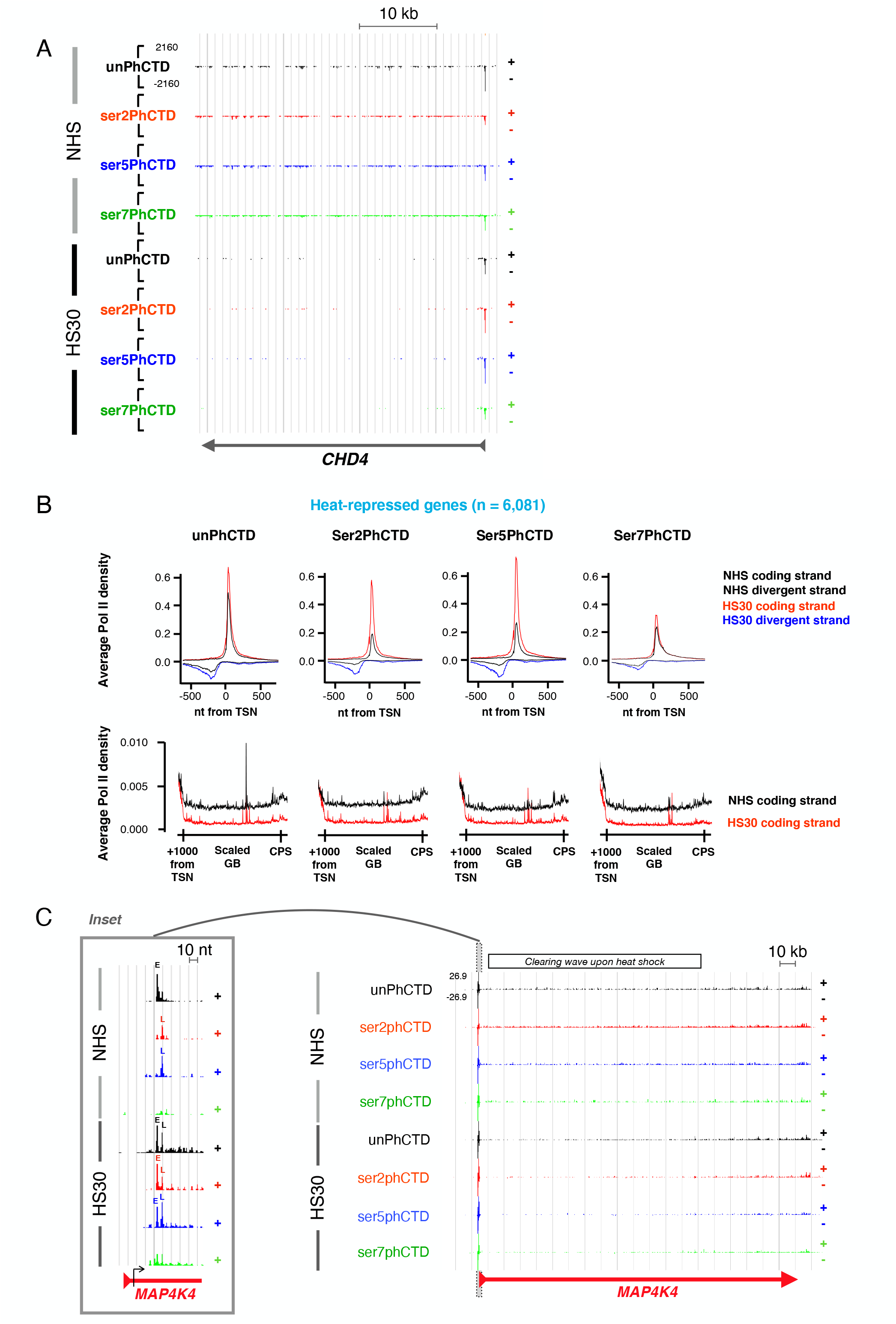
Heat shock captures Pol II with phosphorylated CTD to proximal and distal pauses. **A)** Heat shock represses *CHD4* gene by capturing Pol II to the promoter-proximal region and clearing transcription complexes from the gene body. Please note that the heat shock-triggered increase in promoter-proximal pausing constitutes primarily of serine-2 and serine-5-phospohorylated CTD. **B)** Average PRO-IP-seq densities with indicated Pol II CTD at promoter-proximal regions (upper panels) and gene bodies (lower panels) of heat-repressed genes. The black lines depict PRO-IP-seq signal in nonstressed (NHS) and red (coding) and blue (noncoding) in 30 minute heat shocked (HS30) cells. TSN: Transcription start nucleotide. **C)** Distribution of CTD modifications on *MAP4K4*. Please note that *MAP4K4* is a long gene where the clearing of Pol II from the gene body has not reached the end of the gene during a 30-minute heat shock. The Pol II clearing wave is indicated above the graph. The inset shows distribution of CTD modifications at early (E) and late (L) pauses. The black arrow shows the +1 nucleotide detected from the nascent transcripts.

Despite the massive accumulation of Pol II to promoter-proximal pause at heat-repressed genes, serine-7 phosphorylation at the pause remained minimal (Figure 6A-C). This indicates that even heat-triggered pausing of Pol II at the promoter-proximal region does not stabilize Pol II bearing serine-7-phosphorylation. Thus, serine-7 phosphorylation appears to be a modification intimately connected to the Pol II pause release, and the following assembly of elongation complex that prepares Pol II to its journey through the gene.

**Figure 7.**
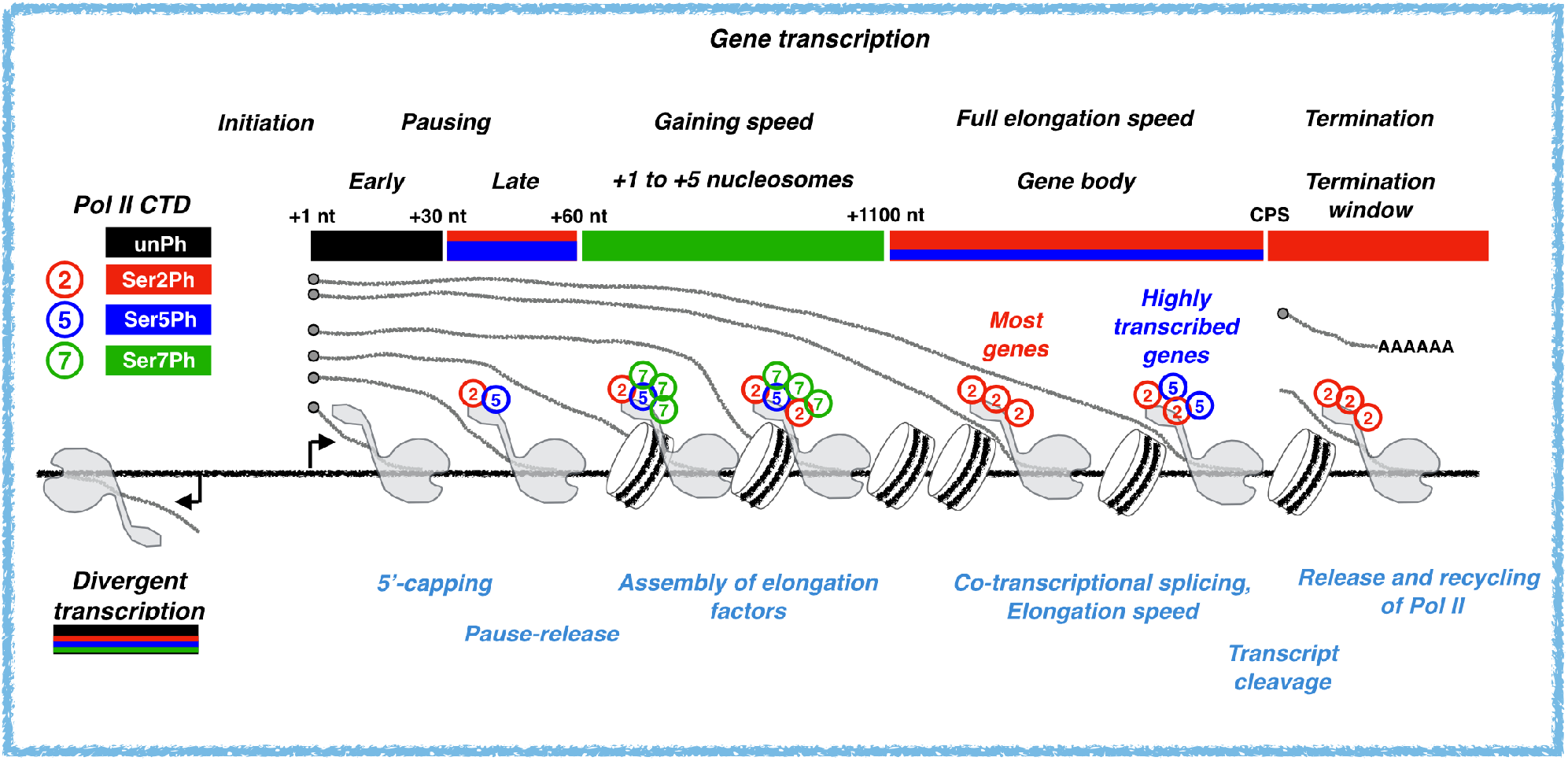
Phosphorylation of Pol II CTD from initiation to transcription termination. **A)** Schematic representation of Pol II CTD phosphorylation status as the transcription proceeds from the initiating nucleotide (+1 nt), through early and late promoter-proximal pause regions, into open chromatin across +1 to +5 nucleosomes, gene body and termination window. The major CTD modifications are indicated with color code above the gene, and co-transcriptional processes occurring at each region listed below the gene.

## Discussion

### PRO-IP-seq provides a nucleotide-resolution tool to discover dynamic changes in engaged Pol II

Pol II is a multi-subunit protein complex whose composition and molecular modifications change as RNA synthesis proceeds through the rate-limiting steps of transcription and transcription coupled RNA processing. Pol II progression at genes and enhancers has been analyzed at nucleotide-resolution (reviewed in Wissink *et al*., 2019), and structures of the PIC, as well as pause and elongation complexes are available in elegant detail *in vitro* (reviewed in Osman and Cramer, 2020). We have lacked efficient tools to monitor nucleotide resolution changes in Pol II complexes *in vivo*. Multiple techniques use immuno-selection of chromatin-bound Pol II, some of which sequence the associated RNAs to track RNA synthesis (reviewed in Wissink *et al*., 2019). These techniques include Pol II ChIP-seq (Hintermair *et al*., 2014) and mammalian NET-seq (mNET-seq; Nojima *et al*., 2015) that have uncovered serine-2-phosphorylation of Pol II CTD to enrich at the 3’-ends of genes while serine-5- and serine-7-phosphorylations dominate at genes’ 5’-regions (reviewed in Zaborovska *et al*., 2016). None of the previous techniques combine nascent transcript labelling and selection of transcription machineries in distinct molecular states to achieve the sensitivity to map the nucleotide-resolution changes in Pol II CTD code. The PRO-IP-seq technique described here 1) labels the active sites of transcription, 2) generates chromatin fragments with a single engaged transcription complex, 3) immuno-selects Pol II by its modifications, 4) purifies the nascent transcripts, 5) maps the active sites of transcription and 6) the precise +1 nts from nascent transcripts, and 7) uses normalization strategy that does not constrain samples to the same total transcript count (Figures 1 and S1-S2). As a result, the precise positional changes in Pol II modifications are mapped from initiation to early and late promoter-proximal pauses and into elongation and transcription termination (Figures 1–3). We expect PRO-IP-seq to be adaptable to also investigate the molecular composition of transcription machineries across the genome, including the dynamic association and disassociation of pausing and elongation factors.

### Nucleotide-resolution CTD code from initiation to promoter escape

*In vitro* studies indicated that Pol II initiates transcription with unphosphorylated CTD (Lu *et al*.,1992). After initiation, Cdk7 phosphorylates serine-5 of the CTD enabling Pol II to break ties with the PIC and the Mediator (Jeronimo and Robert, 2014; Ebmeier *et al*., 2017). Our genome-wide analyses globally corroborate the Pol II that reaches the early pause shortly after initiation has an unphosphorylated CTD (Figures 1–3). Furthermore, Pol II with unphosphorylated CTD peaks at +25 nt and remains high until +30 nt from the initiation. This Pol II at the early pause appears associated with the PIC and has not gone beyond the point of TFIID interaction (Emanuel and Gilmour, 1993; Kwak *et al*., 2013; Fant *et al*., 2020). At the promoter-proximal region, serine-5 phosphorylation enhances 7-methylguanylate capping of the nascent transcript (Komarnitsky *et al*., 2000; Schroeder *et al*., 2000; Nilson *et al*., 2015). The early pause-region has been associated with low, and the late pause-region with high levels of 5’-capping (Rasmussen and Lis, 1993; Tome *et al*., 2018), indicating high-resolution co-occurrence of serine-5-phosphorylation and 5’-capping across the genome (Figure 2).

### What functions as a molecular switch to release the promoter-proximally paused Pol II?

Serine-5, and to a lower extent serine-2, of the CTD was phosphorylated within the pause-region, primarily between +31 to +60 nt from the initiating nucleotide (Figures 1E, 2A and 2C). Since Pol II with unphosphorylated CTD occupies the early pause region (+18 to +31), it’s tempting to speculate that CTD phosphorylation increases the likelihood of Pol II to progress through the pause-region and release to productive elongation. However, provoking a transcriptionally-repressive state at several thousand genes using heat shock revealed that phosphorylation of CTD at serine-2 and serine-5 residues is not sufficient to launch Pol II into elongation (Figure 6). Indeed, the Pol II that accumulated at the pause-regions of heat-repressed genes was abundantly phosphorylated at serines 2 and 5 (Figure 6). These results highlight the need to track the detailed regulatory mechanisms that stabilize Pol II pausing and trigger its release into premature termination or productive elongation. In metazoan species, the release of the Pol II into elongation requires P-TEFb that phosphorylates NELF, SPT5 and serine-2 of Pol II CTD (reviewed in Jonkers and Lis, 2015). Particularly, the CDK9 mediated phosphorylation of SPT5 has been identified as the key trigger to release paused Pol II (Yamada *et al*., 2006). Besides functioning as a signaling platform, CTD phosphorylation has been suggested to aid Pol II escape from a phase-separated domain or condensate around the promoter (Boehning *et al*., 2018). NELF has been reported to form condensates when overexpressed in heat stressed cells, and reduced release of NELF coincides with heat-repression (Rawat *et al*., 2021). Coupling nucleotide-resolution changes in Pol II composition to the progression of RNA synthesis through the rate-limiting steps of transcription should be highly informative in clarifying the regulatory logic of Pol II pause-release, including the functional consequences of P-TEFb, as well as serine-2, NELF and SPT5 phosphorylation.

### Early coding region prepares Pol II to elongation

The median human gene is 20,000 nt long, requiring Pol II to proceed considerable genomic distances to transcribe a pre-mRNA. After being released from the promoter-proximal pause, Pol II proceeds slowly (0.5 kb/min), gaining nearly its full elongation speed (2-5 kb/min) after +1000 nt from the TSS (reviewed in Jonkers and Lis, 2015). We found serine-7-phosphorylation to rapidly increase around the pause-release, peaking at the +1 nucleosome, and remaining high until the end of the MNase-accessible region, +1000 nt from the initiation (Figure 2C). The serine-7-phosphorylation positively correlated with histone acetylation, presence of chromatin modifiers, and intriguingly, predicted transcriptional activity and mRNA abundance (Figures 3 and S5B). Serine-7-phosphorylation has been suggested to prime CTD for recognition by P-TEFb, enabling serine-2-phosphorylation of CTD only at the appropriate stage of the transcription cycle (Czudnochowski *et al*., 2012). These results suggest that distinct kinases could phosphorylate serine-2 of the CTD before and after the pause-release. Indeed, serine-2-phosphorylation was found as a universal mark of active transcription along the gene bodies (Figures 1C and 3A-C). However, serine-5-phosphorylation was enriched along actively transcribed genes in non-stressed cells (Figure 3), and along all genes that escaped the transcriptional repression upon heat shock (Figure S7C). Intriguingly, serine-5-phosphorylation has been coupled to co-transcriptional splicing (Nojima *et al*.,2018), and transcription upon heat shock is known to increase the rate of Pol II elongation to an average 3.5-4.7 kb/min (Mahat *et al*., 2016b; Cugusi *et al*., 2022), suggesting that serine-5-phosphorylation might ensure a fast production of mature mRNA.

In summary, PRO-IP-seq provides a nucleotide-resolution tool to track the molecular modifications of engaged transcription machineries. It uncovered several components of the regulatory logic of Pol II CTD phosphorylation as RNA-synthesis proceeds through rate-limiting steps of transcription. Our study lays the framework to track how Pol II phosphorylation states interplay with its molecular composition and regulation at nucleotide-resolution across the genome.

### Limitations

PRO-IP-seq reports nascent transcripts that associate with transcription machineries in distinct molecular states. The enrichment of Pol II populations in this study is based on antibodies raised against epitopes of Pol II CTD. As with any antibody dependent technique, PRO-IP-seq relies on the specificity of the antibodies. The antibodies used in this study were raised against epitopes containing the same amino acid sequence. For example, the YWG16 antibody was raised against unphosphorylated heptad repeats of Pol II CTD. Although it has been shown to have high specificity to the unphosphorylated Pol II CTD, it likely also detects small amounts of phosphorylated forms of Pol II CTD. This possibility of low level of cross-recognition applies to all the CTD targeting antibodies in this study. To obtain high reliability, we used antibodies that had previously been validated for epitope specificity. Furthermore, for both serine-2 and serine-5 phosphorylation two distinct antibodies were used. Another layer of complexity with Pol II CTD comes from the 52 repeats of T1S2P3T4S5P6S7 present in the human Pol II CTD: Distinct repeats within a single CTD can bear different post-translational modifications. Our study can report the relative levels of each detectable Pol II modifications at nucleotide-resolution across the genome, but it cannot resolve the combinatorial code arising from distinct heptad repeats bearing different serine modifications at a given CTD.

## Supporting information

PRO-IP-seq_Supplemental_Figures

## Acknowledgements

We thank the members of Vihervaara and Lis laboratories for valuable advice and discussions. Alex Greenstone is gratefully acknowledged for the expert assistance with antibody optimization. This work was financially supported by Science for Life Laboratory (A.V.), Swedish Research Council (A.V.), Academy of Finland (A.V.), National Science Foundation Graduate Research Fellowship Program grants DGE-1650441 and DGE-21389899 (P.V.), and NIH grants R01-GM25232 (J.T.L) and RM1-GM139738 (J.T.L.). The content is solely the responsibility of the authors and does not necessarily represent the official views of the NIH. Any opinions, findings, and conclusions or recommendations expressed in this material are those of the author(s) and do not necessarily reflect the views of the National Science Foundation or National Institutes of Health.

## Author contributions

A.V. and J.T.L. conceived and designed the study. A.V. generated the PRO-IP-seq protocol, which was optimized by P.V.. A.V. conducted the computational analyses. All authors interpreted the results and wrote the manuscript.

## STAR Methods

### RESOURCE AVAILABILITY

#### Lead contact

Further information and requests for resources and reagents should be directed and will be fulfilled by the Lead Contact Anniina Vihervaara.

#### Materials availability

This study did not generate new unique reagents.

#### Data and code availability

The PRO-IP-seq datasets generated in this study have been deposited to Gene Expression Omnibus (https://www.ncbi.nlm.nih.gov/geo/), and are available as raw and processed files through accession number GSE200269. Computational analyses have been performed using Unix, R and Python languages. Computational pipelines are available in GitHub: https://github.com/Vihervaara

### EXPERIMENTAL MODEL AND SUBJECT DETAILS

The human K562 erythroleukemia cells were obtained from Lea Sistonen laboratory (Åbo Akademi University, Turku, Finland) and originated from ATCC. The immortalized mouse embryonic fibroblasts (MEFs), which were used for spike-ins, were generated by Ivor Benjamin laboratory (McMillan *et al*., 1998).

### METHOD DETAILS

#### Cell culture and heat shock treatments

Human K562 erythroleukemia cells were cultured in RPMI (Sigma) and the spike-in MEFs in DMEM (Gibco). RPMI and DMEM were supplemented with 10% fetal calf serum, 2 mM L-glutamate, 100 μg ml^-1^ streptomycin, and 100 U ml^-1^ penicillin. Both cell lines were maintained at 37 °C with 5% CO2 and 90% humidity. Both cell lines were tested to be mycoplasma free, and to display morphology, proliferation rate and transcriptional profile characteristic to the respective cell line. The heat shocks in K562 cells were instantly provoked by resuspending 30 million pelleted cells in pre-conditioned pre-warmed media (42°C) and maintaining the cells at 42°C in a water bath for 30 minutes. 30 million non-heat shocked cells were resuspended in 37°C pre-conditioned pre-warmed media and maintained at 37°C for 30 minutes. To avoid provoking transcriptional changes by freshly added media, the cells were expanded 24 h prior to the treatments. The pre-conditioned media was collected from excess cells of the same culture prior to the experiments (Mahat *et al*., 2017).

#### Chromatin isolation

Cells were collected by transferring the cell suspension from 37°C or 42°C into eight times of the volume of ice-cold PBS. The cells were pelleted by centrifugation at 4°C and washed with ice cold PBS. The cell pellets were resuspended in 1 mL ice-cold NUN buffer (0.3M NaCl, 1M Urea, 1% NP-40, 20 mM HEPES pH 7.5, 7.5 mM MgCl_2_, 0.2 mM EDTA, 1 mM DTT), complemented with 20 units SUPERaseIn RNase Inhibitor (Life Technologies # AM2694) and 1× cOmplete EDTA-free Protease Inhibitor Cocktail (Roche cat nr. 11873580001). The samples were vortexed for 30 seconds and centrifuged at 12 500 G at 4°C for 30 minutes. The chromatin pellet (Figure S1A) was washed once in ice cold 50 mM Tris-HCl (pH 7.5) and stored in chromatin storage buffer (50mM Tris-HCl pH 8.0, 25% glycerol, 5mM MgAc2, 0.1mM EDTA, 5mM DTT, 20 units SUPERaseIn RNase Inhibitor) at −80°C.

#### Chromatin fragmentation

Chromatin from 30 million cells was sonicated for 5 minutes at 4°C using Bioruptor (Diagenode) with 30 seconds on / 30 seconds off intervals and high throughput settings. Per each sample of 30 million K562 cells, sonicated chromatin from 250 000 MEFs was added as spike-in material. The same batch of spike-in was utilized for all the samples. Given the smaller genome size of mouse (2.5 x10^9^ bp) compared to human (2.9 x10^9^ bp) and the hypotriploid genome of K562 cells (Naumann *et al*., 2001), the spike-in was estimated to corresponds to 0.5% of total chromatin per run-on reaction. The sonicated chromatin was fragmented to < 80 bp with high concentration RNase-free DNase I for 5 minutes at 37°C water bath [100-300 units of DNase I and 1x DNase I buffer (ThermoFisher, cat.nr. EN0523), 1 mM DTT, 1×cOmplete EDTA-free Protease Inhibitor Cocktail (Roche cat nr. 11873580001) and 20 units SUPERaseIn RNase Inhibitor].

#### Run-on reaction

2x run-on mix [10 mM Tris-Cl pH 8.0, 5 mM MgCl_2_, 1 mM DTT, 300 mM KCl, 50 μM biotin-11-A/C/G/UTP (Perkin Elmer cat nrs. NEL544001EA, NEL542001EA, NEL545001EA, NEL543001EA), 40 units SUPERaseIn RNase Inhibitor, 1% sarcosyl] was prepared and pre-warmed at 37°C water bath. Directly after the 5-minute DNase I treatment, pre-warmed 2x run-on mix was added, and the run-on reaction conducted by maintaining the samples at 37°C for an additional 5 minutes.

#### Immunoprecipitation of Pol II complexes

The run-on reaction was terminated by diluting the chromatin in ice-cold chromatin immunoprecipitation buffer [final concentration 150 mM NaCl, 20 mM Tris-Cl pH 8.0, 1% Triton X-100, 1x proteinase inhibitors, 1x phosphatase inhibitors (phosSTOP, Roche cat. nt. 4906837001), and 20 units of SUPERaseIn RNase inhibitors]. The run-on chromatin was pre-cleared with prewashed protein G beads (Invitrogen Dynabeads, cat. nr. 10004D or GE Healthcare Sepharose, cat nr. GE17-0618-01), and each sample divided into distinct tubes for antibody pulldowns (see Figure 1A). The antibody pulldowns were conducted at 4°C using 5 μg of antibody, pre-bound to 20 μL washed protein G beads, under end-to-end rotation for 1-4 hours. The antibodies used in the pulldown experiments (Figure S1B) were raised against unphosphorylated Pol II CTD (Abcam, YWG16), serine-2-phosphorylated Pol II CTD (Millipore, 04-1571; MBL International, Clone MABI0602), serine-5-phosphorylated Pol II CTD (Millipore, 04-1572; MBL International, Clone MABI0603), and serine-7-phosphorylated CTD (Millipore 04-1570). The negative IgG control antibody was from Santa Cruz (sc-2027). The beads were washed with ChIP wash buffer A (0.1 % SDS, 0.1 % Triton-X 100, 150 mM NaCl, 2mM EDTA pH 8.0, 20 mM Tris-HCl pH 8.0), ChIP wash buffer B (0.1 % SDS, 0.1 % Triton-X 100, 500 mM NaCl, 2mM EDTA pH 8.0, 20 mM Tris-HCl) and ChIP wash buffer C (2mM EDTA pH 8.0, 20 mM Tris-HCl pH 8.0, 10 % glycerol), and diluted in TE-buffer (10 mM Tris-HCl pH 8.0, 1 mM EDTA pH 8.0). NoAb (PRO-seq) control sample was maintained rotating at 4°C without being subjected to antibody pulldowns or antibody-bead washes.

#### Isolation of nascent transcripts

The samples were diluted in TRIzol LS and chloroform, homogenized by vortexing, incubated 1 min on ice and centrifuged 18 000 G, 4°C for 5 minutes. The aqueous layer, containing total RNA in the chromatin-Pol II complexes, was collected, and the RNA pelleted with EtOH using glycoblue (ThermoFisher, AM9516) as a marker of the precipitate. The RNA pellet was washed with 70 % EtOH, air-dried, and diluted in RNase-free water. The RNA was base hydrolyzed with NaOH for 5 minutes and the base hydrolysis terminated with Tris-HCl pH 6.8. NoAb sample was passed through P-30 columns (BioRad, cat.nr. 7326232) to remove free biotinylated nucleotides. The nascent RNAs, containing biotin-nt at the 3’-end, were purified from the non-nascent RNAs using Streptavidin Dynabeads (Invitrogen, MyOne C1, cat. nr. 65002). The bead-nascent-RNA-complexes were washed with PRO-seq wash buffers 1 (50 mM Tri-Cl pH 7.4, 2M NaCl, 0.5 % Triton X-100), 2 (10 mM Tris-Cl pH 7.4, 300 mM NaCl, 0.1 % Triton X-100) and 3 (5 mM Tris-Cl pH 7.4, 0.1), after which the RNA was isolated with TRIzol and chloroform, and precipitated with EtOH.

#### 3’-barcoding and recombining PRO-IP-seq samples

The nascent RNA pellet was diluted in barcoded 3’-adapters, and intermolecular interactions were disrupted at 65°C for 20 seconds. The barcoded adapters were ligated to the nascent transcripts over night at 25°C using T4 RNA Ligase I (NEB, M0204L) in 10 μl reaction volume and 5 μM final 3’-adapter concentration. The sample-specific barcoded 3’-adapters used in this study contain a constant G at the ligation site (5’-end of the adaptor), followed by a 6-nt in-line barcode, and an inverted T at the 3’-end to prevent adapter concatemers (Vihervaara *et al*., 2021). An example 3’-adapter is shown below, indicating the sample-specific in-line barcode inside the square brackets. rG**[rArUrCrArCrG]**rCrGrArUrGrUrGrArUrCrGrUrCrGrGrArCrUrGrUrArGrArArCrUrCrUrGrArArC/3InvdT/ The unligated adapters were removed by binding the nascent transcripts to Streptavidin Dynabeads and washing the beads with PRO-seq wash buffers 1 and 3. During the washes, the samples originating from a run-on reaction were combined in a single new tube (Figure 1A).

#### 5’-decapping, 5’-hydroxyrepair and 5’-adapter ligation

The 5’-decapping and 5’-hydroxyrepair were conducted on-beads. The bead-nascent-RNA-complexes were diluted in RNA 5’ Pyrophosphorylase mix (NEB, M0356S) and incubated at 37°C for 45 minutes. The 5’-hydroxyrepair was subsequently conducted by adding T4 polynucleotide kinase mix (NEB, M0201S) and maintaining the samples at 37°C for additional 45 minutes. The bead-nascent-RNA-complexes were washed with PRO-seq buffers 1 and 3. The nascent transcripts were isolated from the beads with TRIzol and chloroform, and precipitated with EtOH. The air-dried RNA was diluted in 5’-adapter, and the sample incubated at 65°C for 20 seconds. The UMI-containing 5’-adapters were ligated to the nascent transcripts with T4 RNA Ligase I over night at 25°C in 20 μl reaction volume and 2.5 μM final 5’-adapter concentration. The 5’-adapter contained a constant C at the ligation site (3’-end of the adapter), an adjacent 6-nt UMI, and an inverted T at the 5’-end to prevent adapter concatemers (Vihervaara *et al*., 2021). The sequence of the 5’-adapter is shown below, indicating the UMI bolded inside the square brackets.

/5 InvddT/CCTTGGCACCCGAGAATTCCA**[NrNrNrNrNrN]**rC

The unligated adapters were removed by binding the nascent transcripts to Streptavidin Dynabeads and washing the bead-nascent-RNA-complexes with PRO-seq wash buffers 1, 2 and 3. The nascent transcripts were isolated with TRIzol and chloroform, and precipitated with EtOH.

#### Reverse-transcription and amplification

The pellet of nascent transcripts was diluted in a mix of RP1-primer and dNTPs, and intermolecular interactions were disrupted at 65°C for 20 seconds. Reverse transcription was performed with Super Script III reverse transcriptase (ThermoFisher, 18080044) in 25 μl reaction volume containing 2.5 μM RP1-primer, 625 μM dNTPs and 20 units SUPERaseIn RNA inhibitors. The reverse-transcription was conducted using the following settings: 15 minutes at 45°C, 40 minutes at 50°C, 10 minutes at 55°C, 15 minutes at 70°C. The reverse-transcribed samples were test-amplified (as described in Mahat *et al*., 2016a) and visualized on a 6% PAGE. The samples were then amplified in a total of 12 cycles using Phusion Polymerase (in-house) in 1x HF buffer (NEB, M0530S), 1M betaine, 0.25 mM dNTPs, 0.2 μM RP1 primer and 0.2 μM RPI-n index primer. The amplified samples were purified with 1.6x Ampure XP beads (Beckman Coulter, cat. nr. A63880). The samples were analyzed with BioAnalyzer and sequenced with Illumina HiSeq2500 (Cornell University, NY USA) or NovoSeq6000 (NovoGene Inc, CA, USA) using either SE75 or PE150 settings. RP1 and RPI-n primers are Illumina small RNA TruSeq design (Oligonucleotide sequences © 2015 Illumina, Inc. All rights reserved).

#### Computational analyses of PRO-IP-seq datasets

The computational analyses of PRO-IP-seq largely follows the established analyses pipelines of PRO-seq (Kwak *et al*., 2013; Mahat *et al*., 2016a; Vihervaara *et al*., 2021; Rabenius *et al*., 2022). Here, the PRO-IP-seq samples originating from one run-on reaction were barcoded with an in-line hexamer in the 3’-adapter. In the computational analyses, reads in a pool of samples were first separated using fastx barcode splitter (http://hannonlab.cshl.edu/fastx_toolkit/), and when paired-end sequencing was used, correct pairing of reads ensured with fastq_pair (https://github.com/linsalrob/fastq-pair). The adapters sequences (TGGAATTCTCGGGTGCCAAGGAACTCCAGTCAC in Read 1; GATCGTCGGACTGTAGAACTCTGAACGTGTAGATCTCGGTGGTCGCCGTATCATT in read 2) were then removed and the 6-nt UMI in the 5’-adapter was used to collapse PCR duplicates into a single read using fastp (Chen *et al*., 2019). The 7-nucleotide sequences of barcode+C and UMI+C were removed and the Read 1 reverse complemented with fastx toolkit. To avoid mapping human reads to the mouse spike-in genome, we first used high sequencing depth PRO-seq data (80 million uniquely mapping reds) that did not contain foreign DNA (Vihervaara *et al*., 2017), and aligned it to mm10 using Bowtie 2 (Langmead and Saltzberg, 2012). Regions of mm10 where cross-mapping from human nascent transcriptome occurred were masked with maskFasta (bedtools package, Quinlan and hall, 2010). Then, PRO-IP-seq datasets were mapped to the masked mm10 spike-in genome, and uniquely mapped reads used to calculate nf_spike-in_. The PRO-IP-seq reads were subsequently mapped to the human genome (hg19) with Bowtie2. Mapped reads were processed from bam files to bed, bedgraph and bigwig files (Rabenius *et al*., 2022) using samtools (Danecek *et al*., 2021), bedtools (Quinlan and Hall, 2010), and bedgraphToBigWig (https://www.encodeproject.org/software/bedgraphtobigwig/). The complete PRO-IP-seq datasets with raw (fastq) and processed (bigWig) files are available *via* Gene Expression Omnibus database (https://www.ncbi.nlm.nih.gov/geo/) with accession code GSE200269.

#### Normalization of PRO-IP-seq datasets

The samples were normalized using two distinct strategies. First, we generated sequencing-depth-normalized density profiles of the NHS samples, producing patterns of engaged Pol II (Figure S2A) that agree with most of the previously reported results from distinct laboratories (reviewed in Zaborovska *et al*., 2016). This RPM normalization, however, assumes samples to have the same level of total transcription, and it skews analyses when transcriptional activity changes genome-wide, occurring for example during heat shock (Duarte *et al*., 2016; Mahat *et al*., 2016b; Vihervaara *et al*., 2017; 2021). We, therefore, refined the normalization strategy by deriving normalization factors (nfs) from reads mapping to spike-in genome (nf_spike-in_), and to ends of long (>150 kb) genes (nf_longGE_) where heat-triggered changes in transcription had not proceeded during the 30-minute heat shock (Mahat *et al*., 2016b; Vihervaara *et al*., 2017; Booth *et al*., 2018). The nf_spike-in_ was counted against the unphosphorylated CTD from the same run-on reaction (nf_spike-in_ = spike-inCount_sampie_/spike-inCount_unPhCTD_). HS30 sample pool was then adjusted against the NHS pool using the ends of long genes (endCount_unphCTD_HS30_/endCount_unPhCTD_NHS_). In this strategy, a designed control sample (here unPhCTD in NHS condition) gets multiplied with 1. Finally, all the samples are RPM normalized using sequencing depth from the control sample (Figure S2B). This nf-corrected control RPM (nf-cRPM) adjusts samples within an experiment using extrinsic (spike-in genome) and/or intrinsic (unchanged regions) normalization strategies, and then brings transcription profiles in different run-on experiments to comparable y-scale using sequencing depth using the control condition (Figure S2C-D). Importantly, nf-cRPM assumes equal pulldown of spike-in material. Even this strategy is likely to even-out differences between antibodies within a condition, but it does not force the CTD modifications to the same transcriptional range as occurs in traditional RPM normalization.

#### Visualization of genomic data

PRO-IP-seq datasets were visualized using an in-house tool (Hojoong Kwak, Cornell University, Ithaca, USA). The scale of y-axis is linear and equal when comparing signal intensities (tracks) within a browser image. The signal on the plus strand is shown on a positive scale, the signal on the minus strand is shown on a negative scale. The Hi-C data (Rao *et al*., 2014) was visualized using 3D Genome Browser (Wang *et al*., 2018) at 25 kb resolution.

#### Quantification of Pol II densities at genomic loci

The composite profiles of factor intensities from ChIP-seq, enzymatic foot prints from MNase-seq and DNase I-seq, and engaged Pol II complexes from PRO-IP-seq, were generated using bigWig package (https://github.com/andrelmartins/bigWig/). The normalized count of engaged Pol II was queried at defined genomic loci, using a query window of 1 nt, 5 nts, or a 1/50^th^ fraction of the gene body. The queried genomic sites and query windows are indicated in respective figure and figure legend. The average intensities are show with solid lines, the shaded areas display 12.5–87.5% confidence interval.

#### Comparison of phosphorylated Pol II CTD against unphosphorylated CTD

In this study, the antibody raised against unphosphorylated CTD of Pol II (YWG16) was selected for normalization baseline (obtaining nf 1). This antibody was used across all the replicates (Figure S1B), and it was preferred over PRO-seq (NoAb) control due to PRO-seq reporting all nascent transcripts, not only transcription by Pol II. Worth noting is that the protein levels of Pol II, detected with YWG16 antibody, remain unchanged upon and after heat shock exposures in K562 cells and MEFs (Vihervaara *et al*., 2021). The comparison of phosphorylated to unphosphorylated CTD was conducted by first counting the density of engaged Pol II bearing the respective modifications at nucleotide resolution (from normalized bigwig files). Then, the count of engaged Pol II with a given CTD modification was divided with the count of unphosphorylated Pol II CTD at each nucleotide. This ratio was set to log2-scale to center the signal to 0 (see Figure 1C-E).

#### Quantification of transcription and mRNA expression

Transcriptional activity for each gene was measured as previously described (Mahat *et al*., 2016b, Vihervaara *et al*., 2017, Rabenius *et al*., 2022) using gene body as the region marking productive elongation. The normalized gene body count of engaged Pol II molecules was divided with gene body length multiplied with 1000. The resulting gene body RPK (gbRPK) reports the density of Pol II along the gene body. To compare gene transcription in PRO-IP-seq *versus* PRO-seq data, we combined the normalized gbRPK signals from different antibodies in each condition (sum CTD antibodies = unPh + Ser2Ph + Ser5Ph + Ser7Ph). The mRNA expression of respective gene was counted from existing mRNA-seq data in K562 cells (Consortium EP, 2011), remapped to hg19 as previously described (Burchfiel *et al*., 2021).

#### Identification of the Transcription Start Nucleotide (TSN)

The exact transcription start nucleotide was called using the 5’-ends of PRO(-IP)-seq reads as previously described (Vihervaara *et al*., 2021). In brief, the 5’-nucleotide from each read was retained and reported as a bed file. Next, the region −100 to +400 from annotated TSS was queried to find the genomic coordinated with highest count of 5’-nts. The +1 nt was identified from each PRO-IP-seq dataset reported here. Since the +1 nts were highly similar between the PRO-IP-seq datasets (Figure S2), we used +1 nt identified from unphosphorylated CTD under NHS as a representative of the +1 nt (TSN) throughout the study.

#### Identification of the pause nucleotide and pause site distributions

The exact pause nucleotide was called from the 3’-ends of PRO(-IP)-seq reads as previously described (Kwak *et al*., 2013; Vihervaara *et al*., 2021). In brief, the 3’-nucleotide from each read was retained in a bed file. Next, the region −100 to +400 from annotated TSS was queried to find the genomic coordinated with highest count of 3’-nts. The distance of the pause nucleotide from the +1 nt was counted for each gene and plotted as a histogram for the distinct PRO-IP-seq datasets.

#### Western Blotting

Analyses of Pol II protein levels were conducted as previously described (Vihervaara *et al*., 2021) with minor modifications: Cells were lysed in buffer C (25% glycerol, 20 mM HEPES pH 7.4, 1.5 mM MgCl2, 0.42 M NaCl, 0.2 mM EDTA, 0.5 mM PMSF, 0.5 mM DTT), and protein concentration in the soluble fraction was measured using Qubit. 20 μg of total soluble protein was loaded into SDS-PAGE gel and transferred to nitrocellulose membrane (Protran nitrocellulose; Schleicher & Schuell). The membranes were incubated with primary antibodies against Pol II RPB1, either the N-terminal domain (polyclonal, Santa Cruz, sc899), or the CTD (monoclonal antibodies: serine-2-phosphorylated, Millipore, 04-1571; serine-5-phosphorylated, Millipore, 04-1572; and serine-7-phosphorylated, Millipore 04-1570). β-tubulin (Abcam, ab6046) was visualized as a control for loading. The secondary antibodies were HRP conjugated (GE Healthcare), and the blots were developed using an enhanced chemiluminescence method (ECL kit; GE Healthcare). The original scans of the films are available in Mendeley (DOI reserved: 10.17632/s44mkg6jmb.1).

