## Supplementary material for "PRO-IP-seq Tracks Molecular Modifications of Engaged Pol II Complexes at Nucleotide Resolution": PRO-IP-seq_Supplemental_Figures

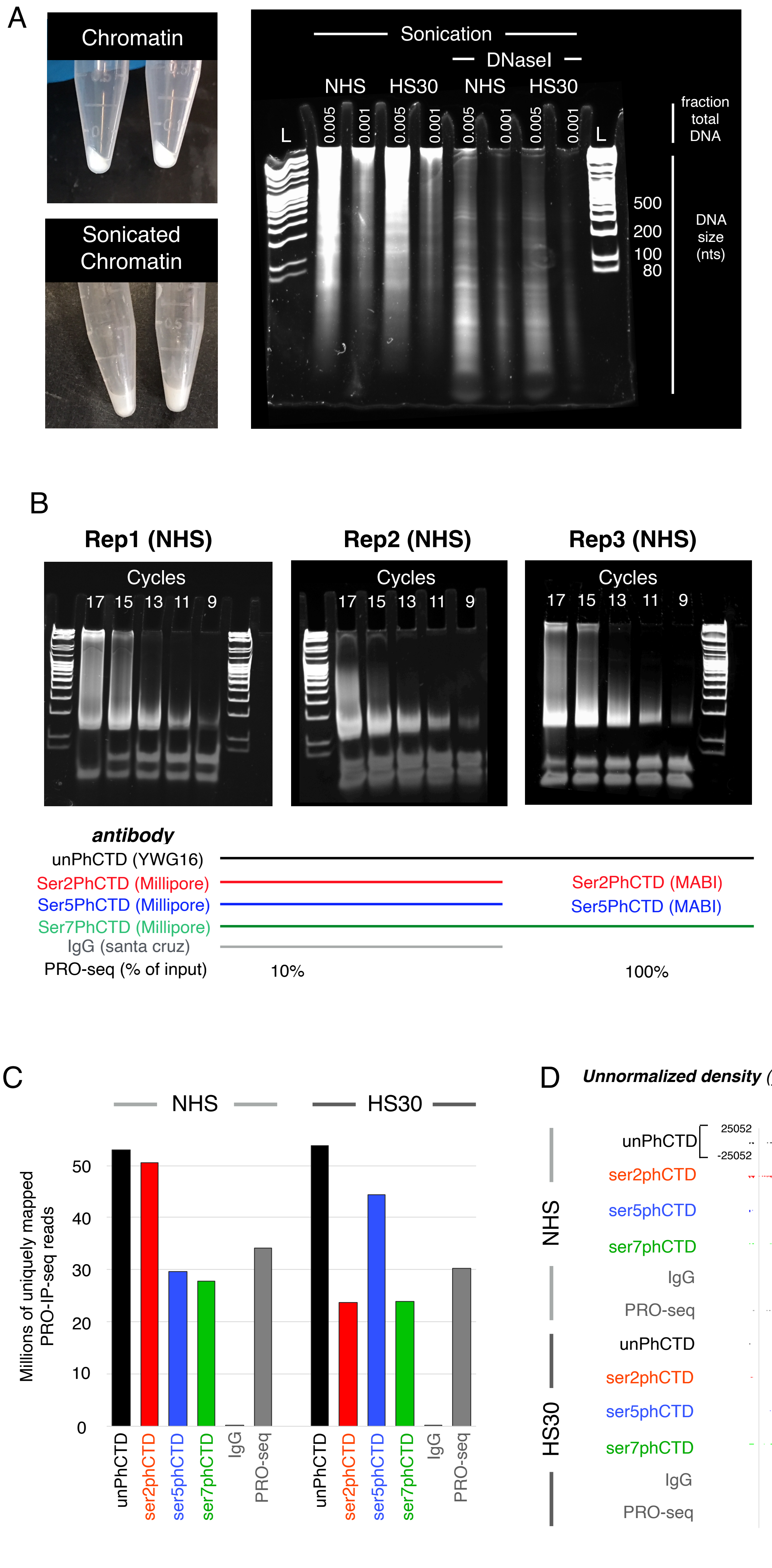


**Figure S1. Generation of PRO-IP-seq libraries. A)** Photographs of chromatin pellets before (*upper left panel*) and after (*lower left panel*) sonication. The chromatin was collected from 30 million of non heat shocked (NHS) or 30min at 42°C heat shocked (HS30) K562 cells. *Right panel*: DNA size distribution before and after DNaseI treatment of the sonicated chromatin. L: ladder. **B)** DNA size distribution in reverse-transcribed and test-ampified PRO-IP-seq samples. Each of the replicates (reps 1-3) contain a pool of barcoded PRO-IP-seq samples indicated below the respective size-distribution. The antibodies were raised against the C-terminal domain of Pol II (CTD) in different states: unphosphorylated (unPhCTD), serine-2-phosphorylated (Ser2PhCTD), serine-5-phosphorylated (Ser5PhCTD), and serine-7-phosphorylated (Ser7PhCTD). IgG serves as a negative control, and no-antibody (PRO-seq) as a positive control. % of input refers to the amount material used in the no-antibody (PRO-seq) positive control, compared to pulldown experiments. **C)** Total count of uniquely mapping reads from the three replicates. **D)** Density profiles of un-normalised PRO-IP-seq samples on an example gene, *RPL23*. The y-scale is indicated in the uppermost browser track (unPhCTD in NHS condition). The y-scale is linear and equal in all the browser tracks. Please note that the negative control (IgG) does not capture nascent transcripts. *RPL23* is a heat-repressed gene in K562 cells, showing promoter-proximal Pol II pausing upon heat shock (Vihervaara et al., 2017).

Supplemental Figure 2

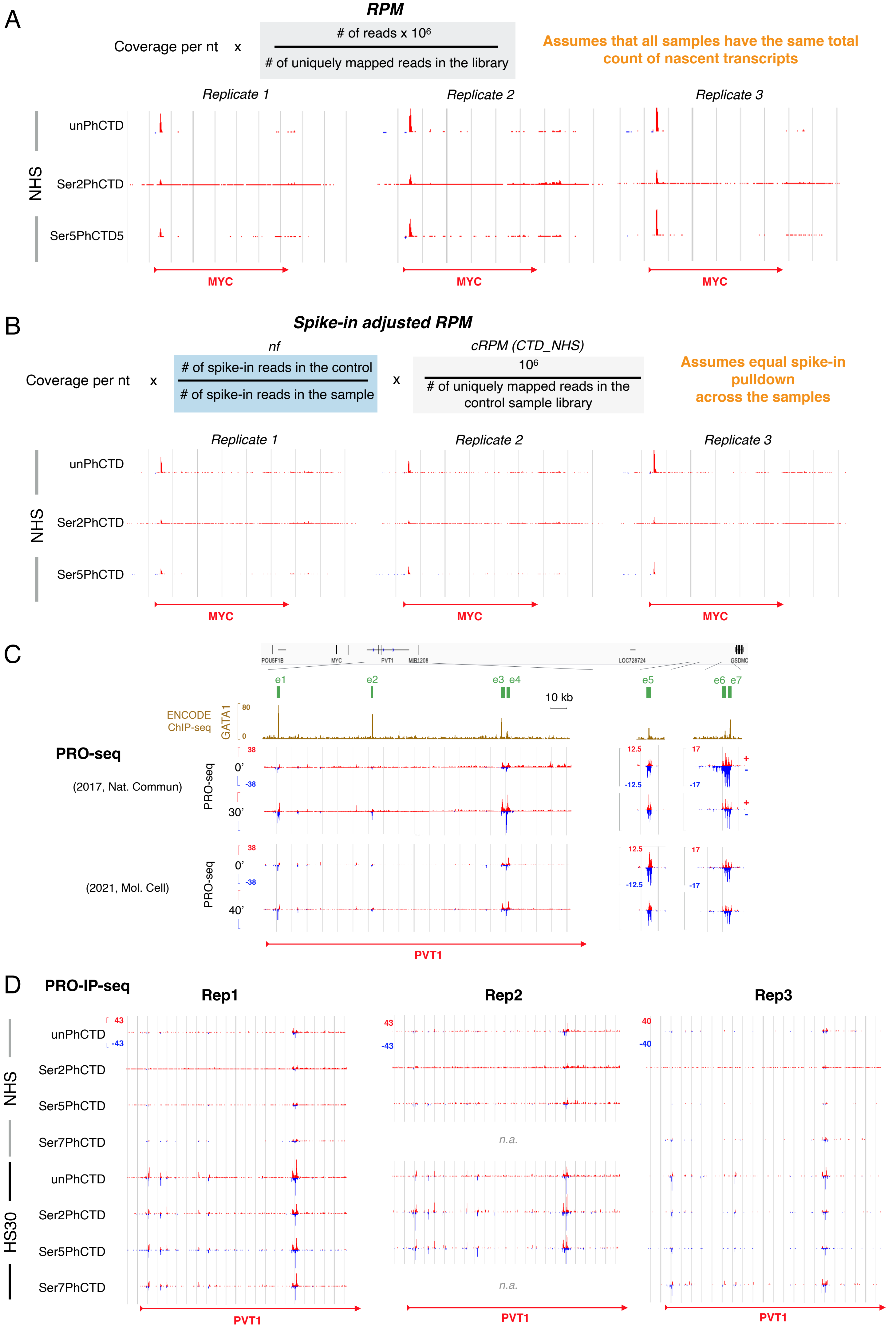

**Figure S2. Normalization of PRO-IP-seq datasets. A-B)** PRO-IP-seq densities displayed on an example, *MYC*, gene after normalization of the samples with **A)** commonly used *reads per millions of uniquely mapped reads (RPM)* normalization, and **B)** *normalization factor (nf) -corrected control RPM (nf-cRPM)* normalization. The shown PRO-IP-seq samples were included in all the three replicates, and are displayed in non heat shock (NHS) conditions. In the *nf-cRPM normalization* **(B)**, the nfs were first derived from the spike-in reads in each replicate, adjusting the samples against the control (unPhCTD) in each replicate. After the nf-normalization, all the samples within a replicate were adjusted for the sequencing depth using total uniquely mapping reads from the control sample. **C-D)** Utilisation of the *nf-cRPM* normalisation strategy brings samples from different experiments and with different sequencing depths to the same y-scale, exemplified with previously published PRO-seq (Vihervaara *et al*., 2017; 2021) experiments **(C)** and with the different PRO-IP-seq replicates **(D)**. In (D), all the replicates and samples used in this study are shown. The regions in C-D cover functionally annotated MYC enhancers (Fulco *et al*., 2016). The coordinates of the enhancers are indicated as green blocks, and binding of a lineage-specific factor in erythrocytes, GATA1 (consortium EP, 2011), is illustrated(golden browser track). Enhancers were chosen as an example for *nf-cRPM* normalization as their activity varies between the cell types, and the detection of enhancer transcription requires highly sensitive strand-specific capture of nascent transcripts. Please also note that enhancers generally gain engaged Pol II upon heat shock as previously observed (Vihervaara *et al*., 2017).

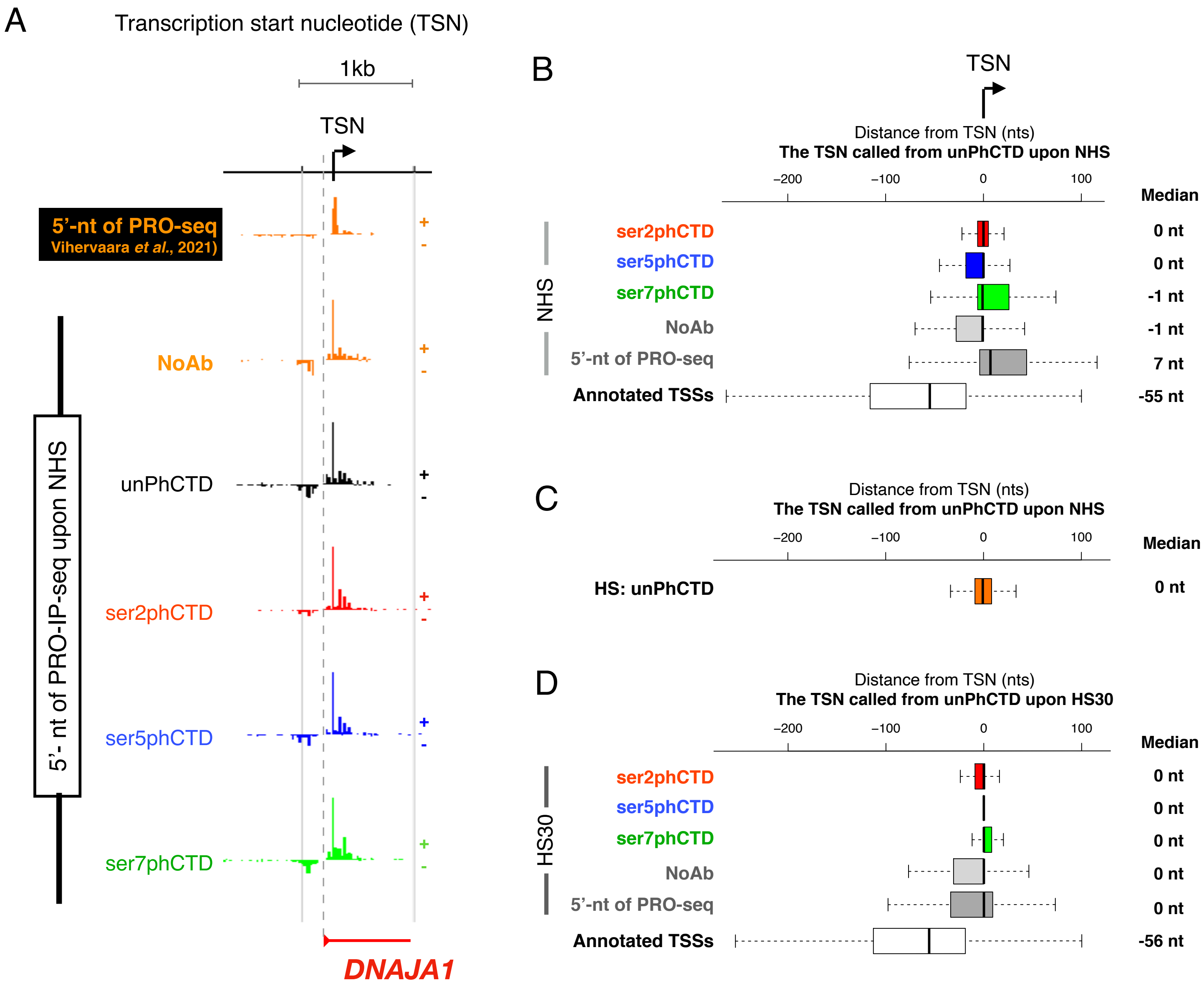


**Figure S3. Transcription start nucleotide (TSN) reports the base (+1) that initiates transcription and is called from 5'-ends of PRO-IP-seq reads. A)** Density profile of 5'-ends of uniquely mapped reads from PRO-seq and PRO-IP-seq experiments, shown on an example gene promoter of *DNAJA1*. The read arrow below the browser tracks shows the start of the annotated gene, and the dashed gray line indicates annotated transcription start site (TSS) from the RefSeq hg19 genome file. The transcription start nucleotide (TSN) is derived as the genomic coordinate with the highest PRO-IP-signal in the unphosphorylated Pol II CTD (unPhCTD) PRO-IP-seq sample (black browser track). **B)** Genomic distance (nfs) of TSNs identified from distinct samples under non-heat shock conditions (NHS), compared against the TSN called from the unPhCTD in NHS condition. The bottommost bar shows the distance of the annotated TSSs (refSeq hg19) to the PRO-IP-seq-identified TSN. **C)** Comparison of TSNs localization under non-heat shock (NHS) and heat shock (HS30) conditions, derived in respective conditions from the unPhCTD PRO-IP-seq sample. **D)** Comparison of TSNs identified from distinct samples under HS30 against the TSN called from the unPhCTD condition HS30. The bottommost bar shows the distance of the annotated TSSs (refSeq hg19) to the PRO-IP-seq-identified TSN.

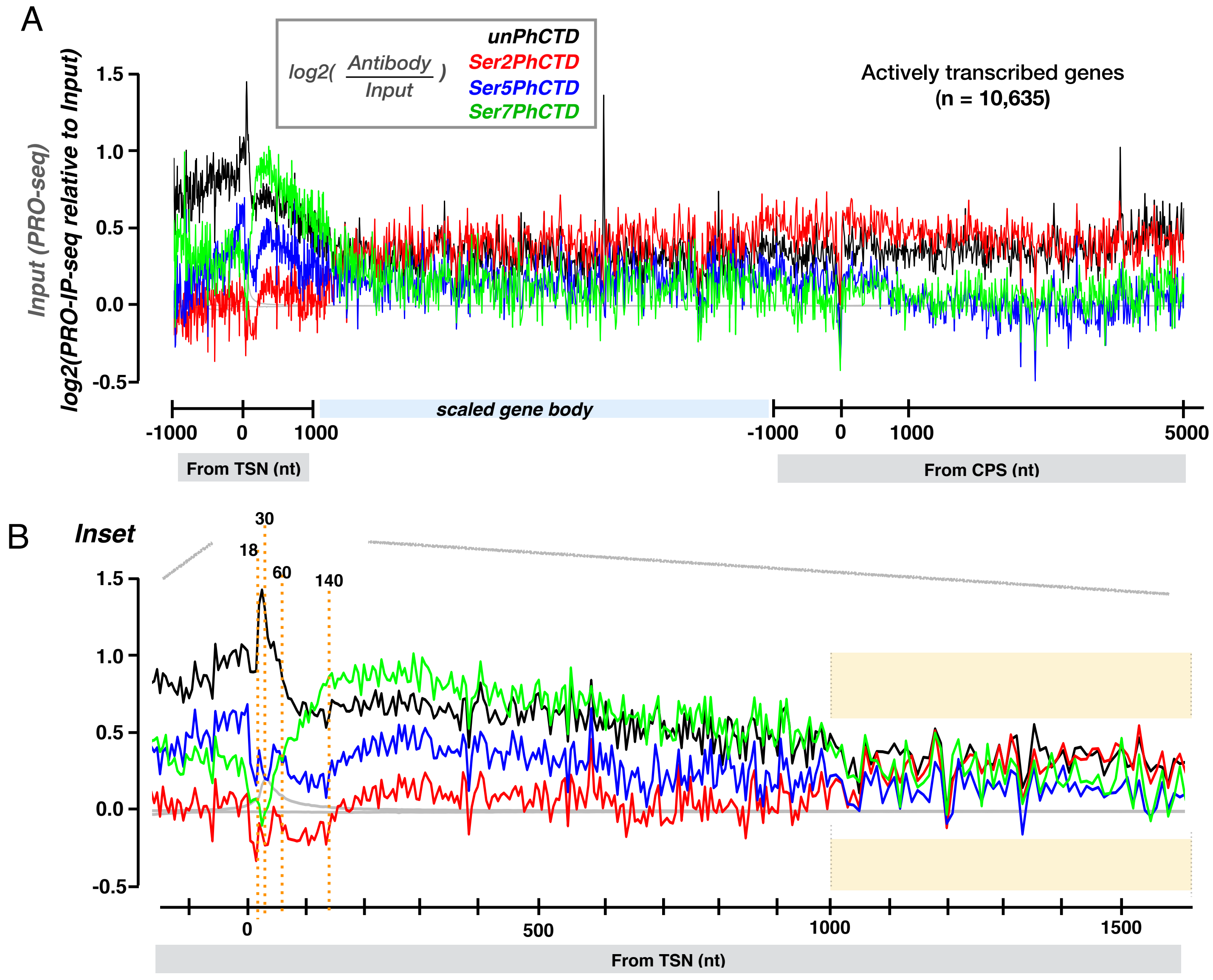


**Supplemental Figure 4. Pol II CTD phosphorylation modifications compared to total transcription. A)** Phosphorylation of Pol II CTD at serines 2, 5 and 7 (PRO-IP-seq) in relation to total transcription (input, PRO-seq). Promoter-proximal region is indicated in a linear scale from -1000 to +1000 nt from transcription start nucleotide (TSN, +1) and is queried in 1-nt windows. Gene body is scaled to 50 bins per gene. Region around CPS and termination window is a linear scale from -1000 to +5000 nt from the CPS. The CTD phosphorylations and total transcription were mapped across all actively transcribed genes in untreated human K562 cells (n=10,635). Average total transcription (input) is indicated in grey. The log2 ratio of PRO-IP-seq to total transcription [log2(antibody/input)] is black for unphosphorylated Pol II CTD and red, blue and green for serine 2, 5 and 7 phosphorylated Pol II CTD, respectively. **B)** Inset comprising the promoter-proximal and early coding regions. The X-axis is the distance from the TSN, queried in 1-nt windows. The orange dotted lines and boxed area indicate the same genomic coordinates as in Figure 1C-E.

Supplemental Figure 5

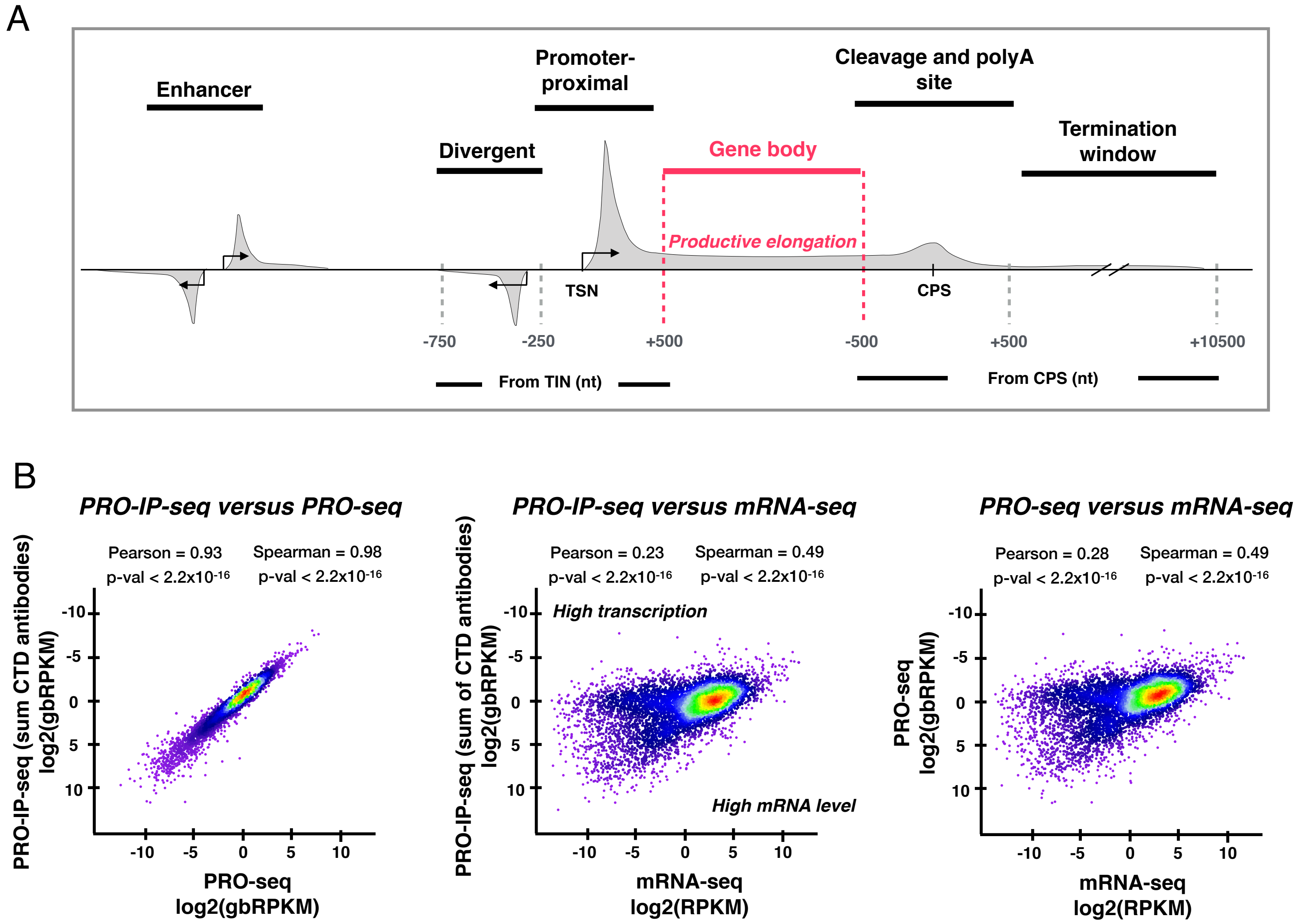


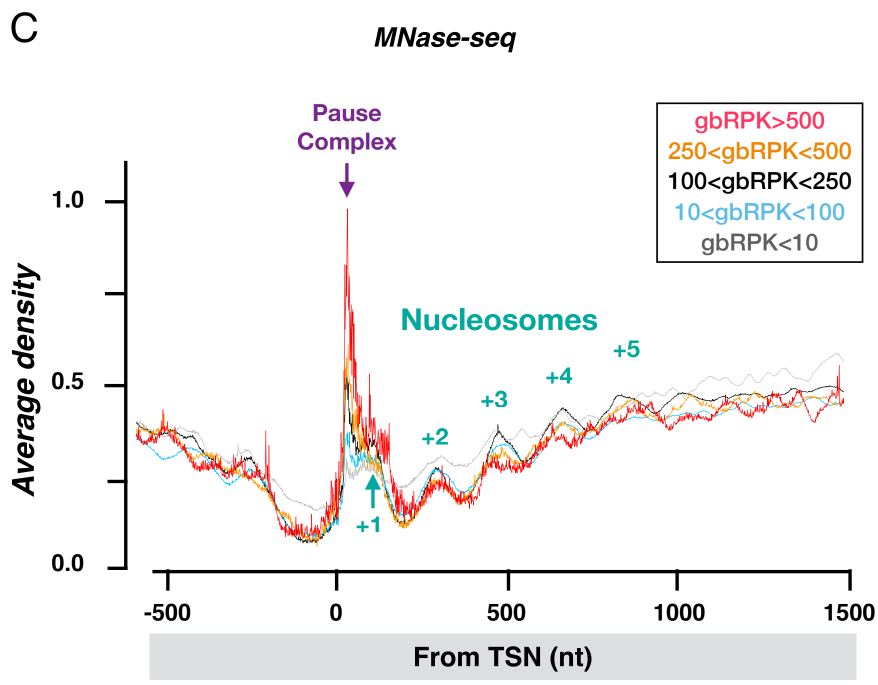


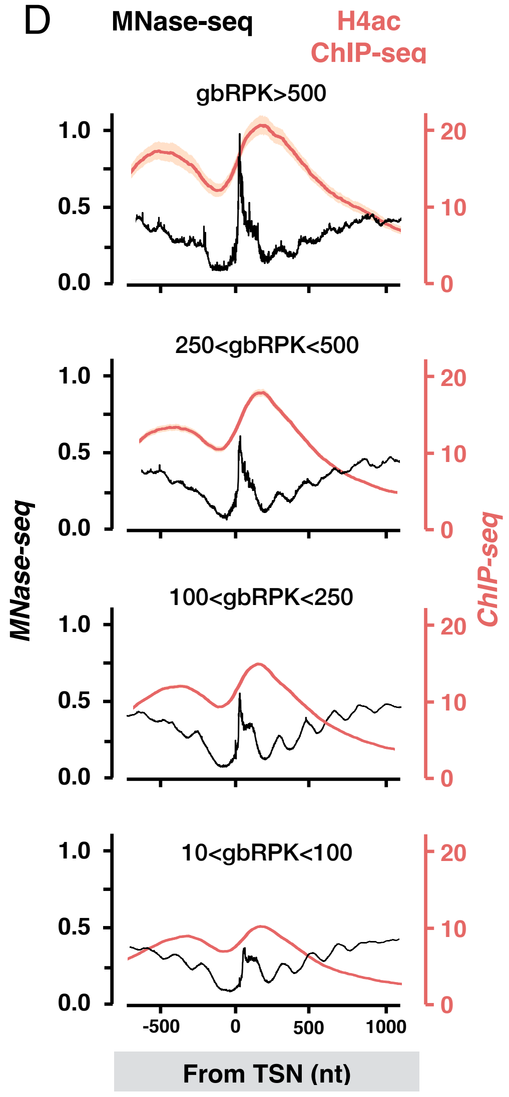


**Figure S5. Correlation of productive elongation with mRNA levels and nucleosomal positions. A)** Schematic illustration of Pol II density across functional genomic regions. The functional genomic regions are as described in Rabenius *et al.*, 2022. Gene body contains Pol II that has proceeded beyond the promoter-proximal pausing and, in this study, is measured as +500 nt from the transcription start nucleotide (TSN) to -500 nt from the cleavage and polyadenylation site (CPS). **B)** Productive elongation from PRO-IP-seq (this study) correlates with productive elongation from Precision Run-On sequencing (Vihervaara *et al.*, 2017) and mRNA expression (Consortium EP, 2011). **C)** Average profile of MNase-seq at the indicated gene groups. The shielded pause complex, and +1 to +5 nucleotides are indicated. **D)** MNase-seq and histone 4 acetylation (H4ac) ChIP-seq at the promoter-proximal region of the indicated genes.

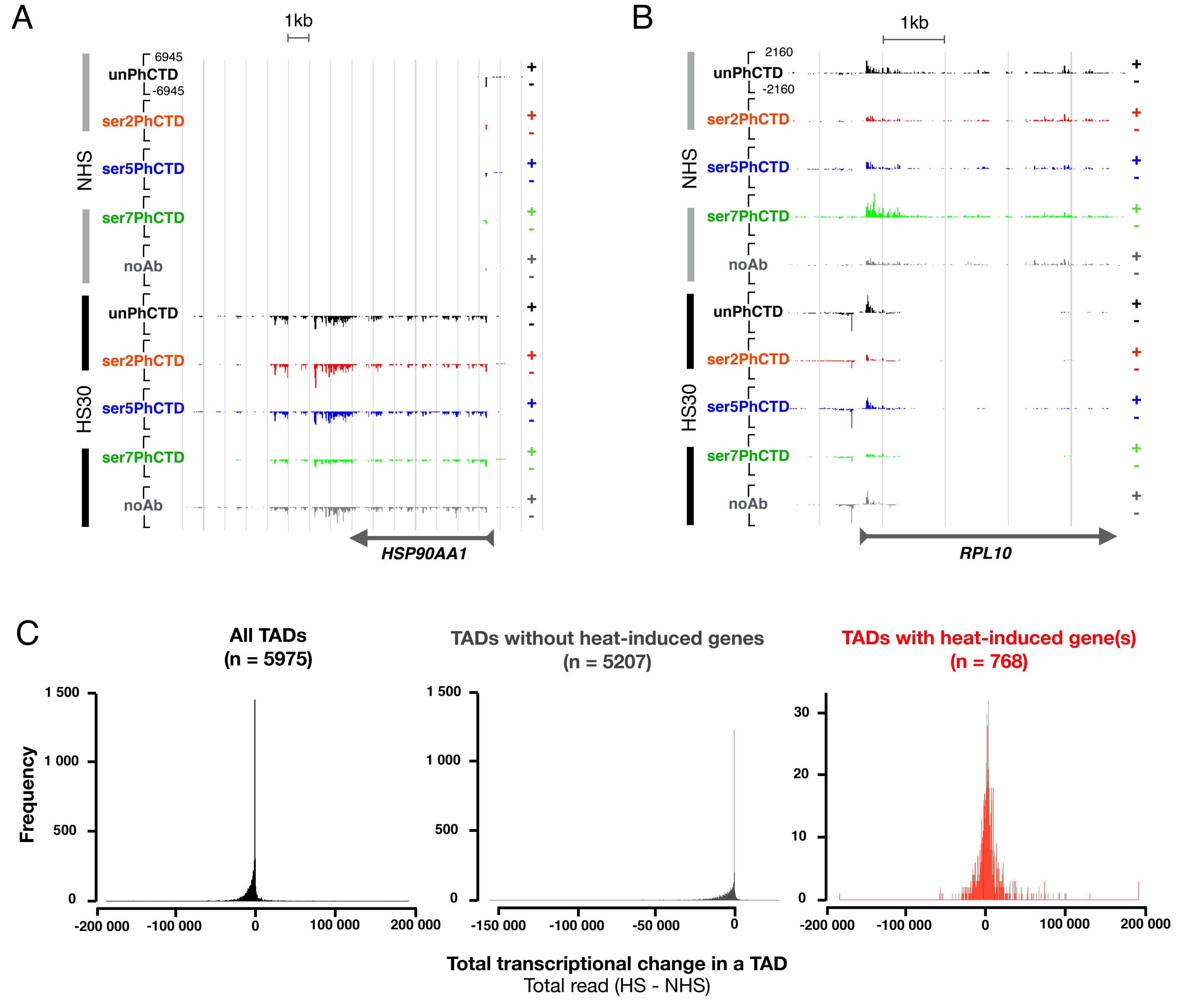


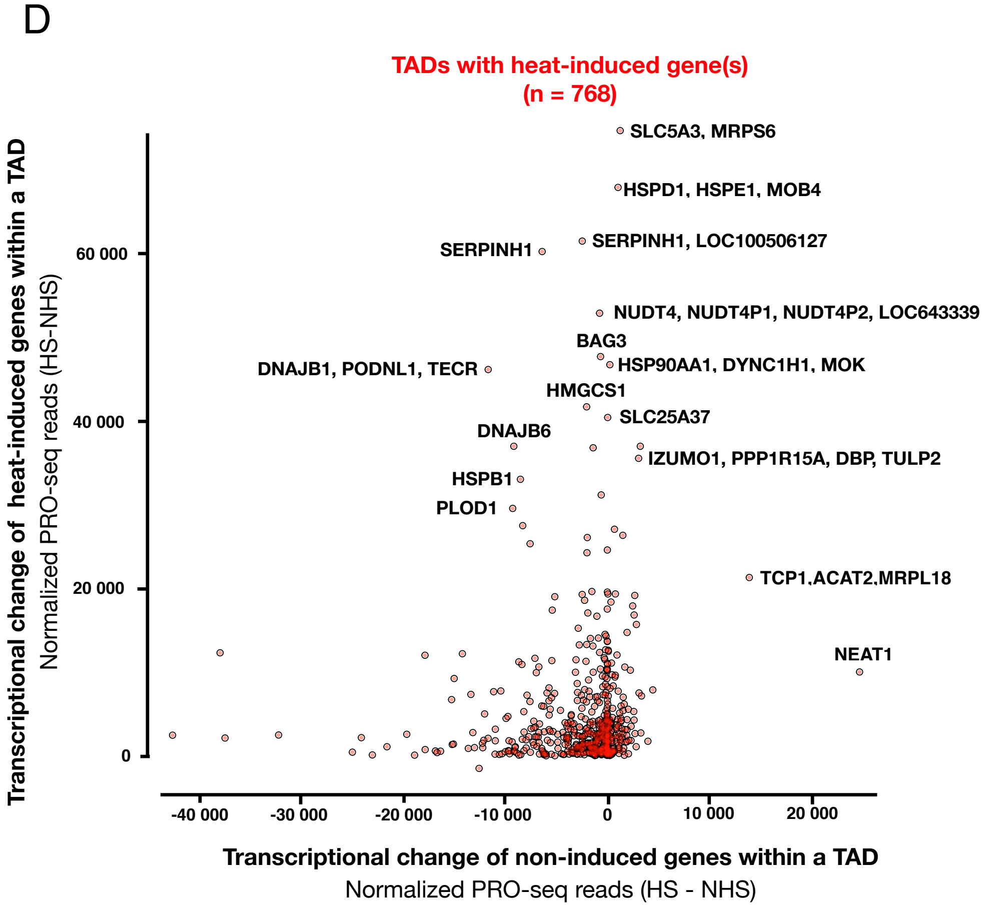


**Figure S6. Transcriptional change in TADs A)** Heat-induced wave of transcription at *HSP90AA1* gene. **B)** Heat-triggered repression at *RPL10* gene. **C** Histograms showing total transcriptional change in TADs. The TADs are as described in Rao *et al.*, 2018. **D)** Transcriptional change at heat-induced genes (y-axis) compared with transcriptional change of non-induced genes (x-axis) in the same TAD (semi-transparent red sphere). The heat-induced genes in selected TADs are indicated. Please note that transcriptional repression in a TAD does not compensate for the increased recruitment of Pol II to the heat-induced genes within the same TAD.

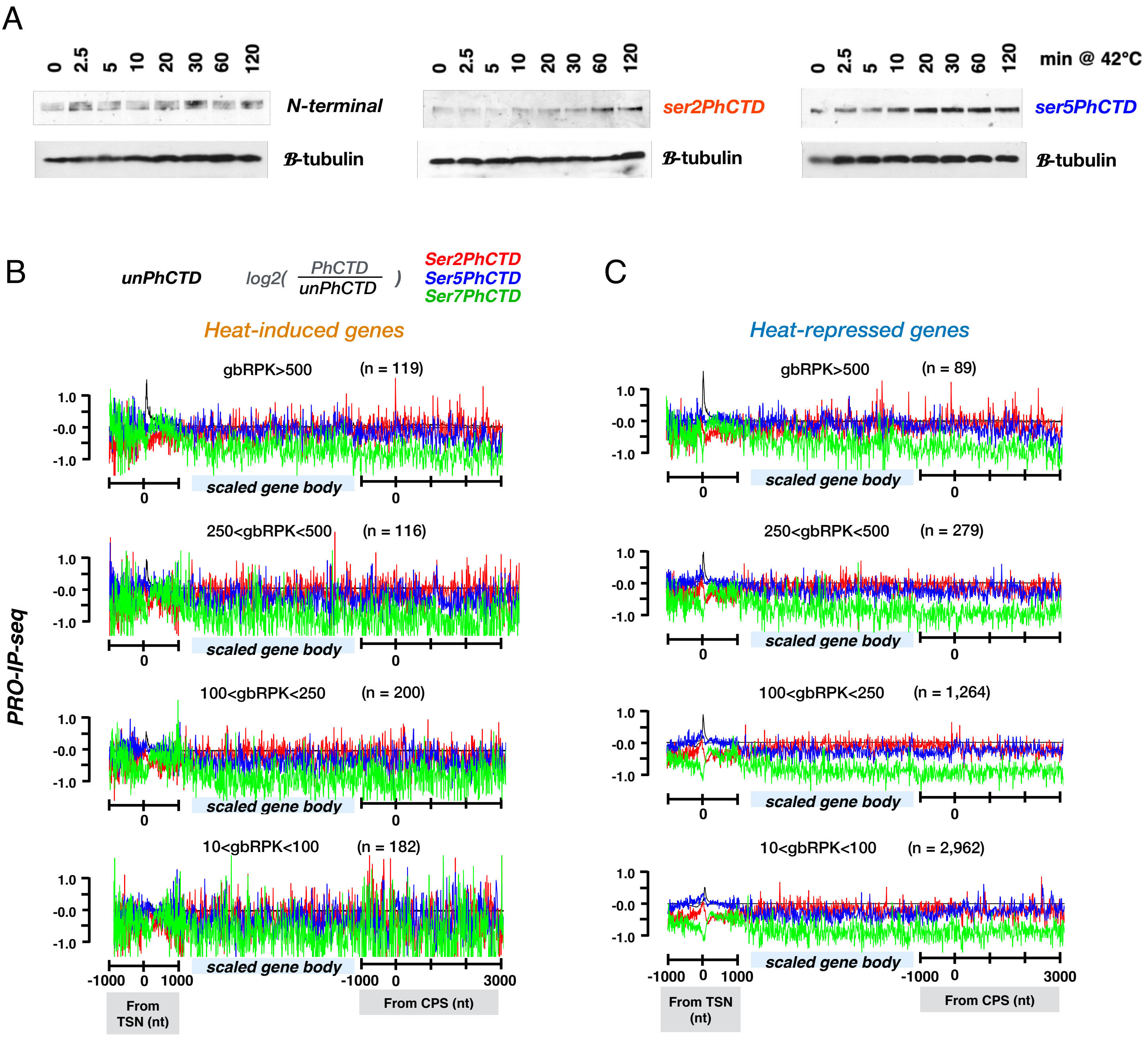


**Figure S7. Heat shock increases serine-5-phohosphorylation at Pol II CTD. A)** Westen blotting detection of Pol Ii and its CTD phosphorylation during a heat shock time course. **B-C)** Relative enrichments of Pol II CTD phosphorylations against unphosphorylated Pol II CTD at **B)** heat-induced and **C)** heat-repressed genes. Each tick in B-C) indicates 1000 nt linear distance from the transcription start nucleotide (TSN).
